# The phylogenetic functional conservation of *Drosophila* Seven-In-Absentia (SINA) E3 ligase and its two human paralogs, SIAH1 and SIAH2, in *Drosophila* eye development

**DOI:** 10.1101/2020.04.28.067074

**Authors:** Robert E. Van Sciver, Yajun Cao, Amy H. Tang

**Author notes:** To whom correspondence should be addressed: Amy H. Tang, Ph.D. Professor of Cancer Biology, Eastern Virginia Medical School, 651 Colley Avenue, Norfolk, Virginia 23501. These two authors share co-1^st^ authorship with equal contribution (Robert E. Van Sciver, Yajun Cao).

## Abstract

Seven-IN-Absentia (SINA) is the most downstream signaling gatekeeper identified thus far in the RAS/EGFR pathway that controls photoreceptor cell fate determination in *Drosophila*. Underscoring the central importance of SINA is its phylogenetic conservation in metazoans, with over 83% amino acid identities shared between *Drosophila* SINA and human SINA homologs (SIAHs). SIAH is a major tumor vulnerability in multidrug-resistant and incurable cancer. SIAH inhibition is an effective strategy to shut down the tumor-driving K-RAS/EGFR/HER2 pathway activation that promotes malignant tumor growth and metastatic dissemination. To further delineate the SINA function in the RAS/EGFR pathway, a genetic modifier screen was conducted, and 28 new *sina* mutant alleles were isolated via ethyl methanesulfonate (EMS) and X-ray mutagenesis. Among them, 26 of the new *sina* mutants are embryonic, larval, or pupal lethal, and stronger than the five published *sina* mutants (*sina*^1^, *sina*^2^, *sina*^3^, *sina*^4^, and *sina*^5^) which are early adult lethal. By sequencing the SINA-coding region of *sina*^ES10^, *sina*^ES26^, *sina*^ES79^, and *sina*^ES473^ homozygous mutant animals, we identified three invariable amino acid residues in SINA’s RING-domain whose single point mutation ablates SINA function. To demonstrate the functional conservation of this medically important family of RING domain E3 ligases in *Drosophila*, we established a collection of transgenic lines, expressing either wild type (WT) or proteolysis-deficient (PD) SINA/SIAH inhibitors of *Drosophila* SINA^WT/PD^ and human SIAH1^WT/PD^/2^WT/PD^ under tissue-specific GAL4-drivers in *Drosophila* eye, wing, and salary gland. Our results showed that *Drosophila* SINA and human SIAH1/2 are functionally conserved. Our bioengineered SINA^PD^/SIAH^PD^ inhibitors are effective in blocking the RAS-dependent neuronal cell fate determination in *Drosophila*.

## Introduction

The RAS signaling pathway plays a pivotal role in controlling fundamental cellular and developmental processes, including cell proliferation, differentiation, motility, and apoptosis [1-4]. Oncogenic K-RAS mutations are detected in approximately 30% of human cancers [4-9]. Persistent K-RAS/EGFR/HER2 pathway activation is prevalent in chemo-resistant, relapsed and metastatic human cancer [2, 10-13]. The clinical reality is that oncogenic K-RAS-driven malignant tumors remain “undruggable,” despite the recent development of small molecule inhibitors that covalently and irreversibly bind to a mutant cysteine in hyperactive K-RAS^G12C^ [14-20]. Although these high-affinity cysteine-targeting covalent K-RAS^G12C^ inhibitors have shown promising antitumor efficacy in phase I/II studies [21-23], their clinical efficacy remains to be demonstrated.

*Drosophila* eye development is a well-established, powerful, and genetically-tractable model system in which to study RAS pathway activation. This elegant system was instrumental in identifying the key signaling components and elucidating the conserved operational principles of the RAS signal transduction cascade *in vivo* [24-28]. Due to the extraordinary degree of evolutionary conservation of RAS signaling pathway in metazoans, the molecular insights and core principles learned from *Drosophila* RAS studies have informed and guided many aspects of mammalian K-RAS studies in human cancer [28-32]. Seven-IN-Absentia (SINA) is an evolutionarily conserved RING-domain E3 ubiquitin ligase that serves as the most downstream signaling hub whose function is indispensable for proper RAS signal transduction [25, 31, 33, 34]. Even in the context of constitutively activated RAS^V12^ or any other hyperactive upstream signals emitted from EGFR/RAS/RAF/MEK/ERK/Pointed/PHYL pathway, the absence of SINA activity completely blocked the R7 photoreceptor cell fate determination, demonstrating that SINA is indeed the most downstream signaling gatekeeper essential for proper RAS/EGFR/MEK/ERK signal transduction in *Drosophila* eye development [24, 25, 33-37].

Guided by ample evidence in developmental, evolutionary, and cancer biology, we proposed an alternative strategy to shutdown “undruggable” oncogenic K-RAS activation by targeting human SINA homologs (SIAHs) in human cancer cells [28-32, 34]. Using a proteolytic deficient (PD) SIAH mutant to block SIAH1/2 function, we demonstrated that human SIAHs are a major tumor vulnerability of oncogenic K-RAS/EGFR-driven tumorigenesis and metastasis in human pancreatic and lung cancer [29, 30] (triple negative breast cancer data not shown). Hence, SIAH is strategically well-positioned to become a new, actionable, and potent anti-K-RAS drug target to eradicate chemo-resistant, incurable, and metastatic human cancers [28-30, 32, 38, 39].

The central importance of SINA/SIAH biology in RAS pathway activation lies in the extraordinary degree of evolutionary conservation of these RING-domain E3 ligases across metazoan species [31]. There are over 83% amino acid identities shared between *Drosophila* SINA and its human homologs (SIAHs) [31, 40]. To date, there are five published *sina* mutants that are hypomorphic alleles [33]. The strongest among them, *sina*^2^ and *sina*^3^ mutants, resulted in truncated SINA proteins missing one-third (32.5% and 33.1%, respectively) of the C-terminal sequence [33]. Hence, there was a clear need to conduct unbiased mutagenesis screen to isolate and identify new *sina* mutant alleles to reveal the key amino acids and core functional domains indispensable for proper SINA function. New genetic studies are needed to compare and delineate the evolutionarily conserved function of *Drosophila* SINA and human SIAH in the RAS signaling pathway in a cross-species study [31, 33, 34].

In this study, we compared and delineated the functional conservation of this medicinally important and evolutionarily conserved family of SINA/SIAH RING-domain E3 ligases in *Drosophila* R7 photoreceptor development. Firstly, we conducted a saturated *GMR-phyl* modifier screen and isolated 28 new *sina* mutant alleles by ethyl methanesulfonate (EMS) and X-ray mutagenesis. By sequencing the *sina*^EMS^ mRNA transcripts from *sina*^ES10^, *sina*^ES26^, *sina*^ES79^, and *sina*^ES473^ homozygous viable mutant animals, we identified three invariable amino acids in SINA’s zinc-coordinating RING-domain whose single point mutation ablates SINA function, supporting our SIAH^PD^ inhibition strategy. Secondly, we compared the functional conservation of *Drosophila* SINA and human SIAH across species *in vivo*. We found that these phylogenetically conserved *Drosophila* SINA and human SIAH1/2 are likely to be functionally interchangeable. We reported that SINA^PD^/SIAH^PD^ inhibitors can effectively block the RAS-dependent neuronal cell fate determination in *Drosophila.*

## Results

### Historic Perspective: The five pre-existing *sina* mutant alleles are hypomorphic alleles

By conducting an F2 genetic screen, five *sina* alleles (*sina*^1^, *sina*^2^, *sina*^3^, *sina*^4^, and *sina*^5^) were isolated, that revealed critical SINA function in *Drosophila* sensory neuron cell fate determination [33]. A major limitation of this labor-intensive F2 screen was the hidden prerequisite that homozygous *sina* mutant animals had to survive to adulthood in order to detect the missing R7 photoreceptor phenotypes by pseudo-pupil assays [33]. Thus, any embryonic, larval or pupal lethal *sina* mutant alleles could not be identified from such an F2 genetic screen [33]. Genetic complementation tests described below indicated that these five previously published *sina* mutant alleles were not null alleles, but rather hypomorphic alleles.

It was reported that homozygous *sina*^1^, *sina*^2^, *sina*^3^, *sina*^4^, and *sina*^5^ mutant animals were viable as young adults who exhibited PNS mutant phenotypes including the missing R7 photoreceptor cells, missing sensory bristles (e.g. macrochaetes and microchaetes), reduced adult lifespan, reduced fertility and overall survival [33]. Both *sina*^2^ and *sina*^3^ mutant alleles showed a premature stop codon at amino acid residues #210 or #212, respectively, resulting in a C-terminal-truncated SINA protein that is missing one-third (32.5% or 33.1%) of its coding sequence, including the SINA dimerization domain [31, 33]. This large deletion provided little insight into critical amino acids that could guide precision drug design of SIAH inhibitors against oncogenic K-RAS-driven malignant tumors in the future. Hence, there was a clear need to isolate and identify strong *sina* point mutations and null alleles to study the key amino acids and core functional domains critical for SINA function in RAS-dependent photoreceptor cell fate determination in *Drosophila*.

### 28 new *sina* mutant alleles were identified from *GMR-phyl* genetic modifier screen

Phyllopod (PHYL) is a RAS/RAF/MEK/MAPK-induced protein that functions directly upstream of SINA [35, 36]. PHYL is a known SINA-interacting protein [34]. Mechanistically, PHYL recruits SINA (an E3 ligase) and EBI (an F-box protein) to target TTK^88^, a transcription factor and neuronal repressor, for protein degradation to unleash RAS-driven neuronal development in R7 photoreceptor cells [34]. By overexpressing PHYL in *Drosophila* eye, we aimed to isolate new *sina* mutant alleles by conducting an F1 genetic modifier screen, and we anticipated that a loss of SINA function should suppress *GMR-phyl* rough eye phenotypes *in vivo*.

To conduct an unbiased genetic screen to isolate new *sina* point mutations and other strong null alleles, we performed a saturated *GMR-phyl* genetic modifier F1 screen using both a low dose ethyl methanesulfonate (EMS) and X-ray radiation mutagenesis, and we established 8 independent enhancer groups and 11 independent suppressor groups (*GMR-phyl* genetic screen, Tang lab manuscript in preparation). The *sina*^loss of function^ modifier group was the largest suppressor group and it was also the strongest suppressor of *GMR-phyl* rough eye phenotype. From this F1 modifier screen, we successfully isolated 28 new *sina* mutant alleles, including 11 EMS and 17 X-ray *sina* mutant alleles in support of our SINA function study.

Complementation tests of these 28 *sina* mutant alleles were conducted with (1) a large *sina* genomic deletion allele – DF(81K72) [33], (2) *sina*^2^, (3) *sina*^3^, and (4) with each other. These elaborate genetic results confirmed that these newly identified 28 *sina* mutant alleles are indeed *sina*^loss of function^ mutant alleles. Importantly, all 28 new *sina* mutant alleles are stronger than the five published *sina* hypomorphic mutant alleles, namely *sina*^1^, *sina*^2^, *sina*^3^, *sina*^4^, and *sina*^5^, based on the suppression of *GMR-phyl* rough eye phenotype, as well as the early homozygous lethality as shown in complementation tests. Among 28 new *sina* mutant alleles, 26 of them are homozygous lethal at embryonic, larval, or pupal stages prior to eclosion (**Table 1**). Only a small fraction of these two homozygous mutant animals (*sina*^ES26^ and *sina*^ES10^) can survive to adulthood for a few hours after eclosion. 11 EMS-induced *sina* mutant alleles completely suppressed the rough eye phenotype of ectopic *GMR-phyl* expression in *Drosophila*. These 11 *sina*^EMS^ mutant alleles exhibited stronger mutant phenotypes than those of the published C-terminal truncated *sina*^2^ and *sina*^3^ alleles. By conducting a *GMR-phyl* F1 modifier screen as designed, we successfully isolated 28 new *sina* mutant alleles from this genetic screen. Based on complementation tests, we report that *sina* null alleles are embryonic lethal. *Sina* is an essential gene in normal development, and the complete loss of SINA function is cell lethal and embryonic lethal (**Supplemental Figure S1**).

**Table 1.** Summary of *sina* mutant alleles.

| ***sina* allele** | **Nucleotide Change (ORF)** | **Amino Acid Change** | **Domain** |
| --- | --- | --- | --- |
| *sina*<sup>ES10</sup> | G (311) → A | Cys (104) → Y | RING |
| *sina*<sup>ES26</sup> | 10 bp deletion 435-444 | E145 Frameshift mutation<br>Change in 25 amino acids<br>and early truncation | SIAH-Type Zinc<br>Finger |
| *sina*<sup>ES79</sup> | G (218) → A | Cys (73) → Y | RING |
| *sina*<sup>ES473</sup> | G (227) → A | Cys (76) → Y | RING |
| **Previously Published Alleles [33]** |  |  |  |
| *sina*<sup>1</sup> | T (728) → G | Leu (243) → R | Between SBS and<br>DIMER |
| *sina*<sup>2</sup> | G (630) → A | Trp (210) → Stop | SBS |
| *sina*<sup>3</sup> | GGT (630) Δ GGT (939) | Met (212) → Stop | Between SBS and<br>DIMER |
| *sina*<sup>4</sup> | C (550) → T | His (184) → Tyr | SZF |
| *sina*<sup>5</sup> | C (700) → T | His (234) → Tyr | Between SBS and<br>DIMER |

### Three new *sina*^EMS^ point mutations in the RING domain exhibit strong mutant phenotypes

We first focused on 11 EMS-induced *sina* mutant alleles that completely suppressed the rough eye phenotype of ectopic *GMR-phyl* transgene expression (Tang lab manuscript in preparation). We found that these 11 new *sina*^EMS^ mutant alleles exhibited homozygous mutant lethality at embryonic, larval, pupal, or early adult stage. To overcome and bypass the homozygous lethality of these 11 new *sina*^*EMS*^ mutant alleles, we crossed each of the early homozygous-lethal *sina* mutant alleles, *sina*^*EMS*^*/TM6B* flies, individually to that of a hypomorphic *sina*^*3*^ mutant allele, *sina*^3^*/TM6B* flies, to generate a viable heterozygous *sina*^*EMS*^/*sina*^*3*^ mutant animals with an intermediate mutant phenotype. These resultant genetic crosses generated a number of heterozygous long larvae, some long pupae, and/or a few adults that eclosed and died as young adults. From these eleven heterozygous *sina*^*EMS*^/*sina*^*3*^ mutant animals, we generated cDNA libraries, conducted *sina* open reading frame (ORF)-specific RT-PCR amplification, and sequenced four *sina*^*EMS*^ mutant alleles (**Figures 1A and 1B**). By strategically positioning the 3’ RT-PCR primer in the deleted C-terminal region of the *sina*^3^ allele, only the mutant *sina*^EMS^ coding sequences were selectively amplified. By using this *sina* ORF-specific RT-PCR scheme, the EMS-induced point mutation(s) in the *sina*^EMS^ mutant ORFs were sequenced and validated independently (**Figures 1A and 1B**). The RT-PCR results revealed that 4 of the 11 *sina*^*EMS*^/*sina*^*3*^ heterozygous long larvae generated full-length *sina*^*EMS*^ ORF transcripts, demonstrating the four *sina*^EMS^ mutant alleles (*sina*^ES10^, *sina*^ES26^, *sina*^ES79^ and *sina*^ES473^) harbored valuable point mutations or a small truncation or deletion (**Figure 1B**). High-fidelity RT-RCR were performed by using three-independent mRNA isolates to generate three-independent cDNA libraries prepared from three separate batches of heterozygous *sina*^*EMS*^/*sina*^*3*^ mutant larvae, and identical point mutations were validated and confirmed successfully (**Figure 1A**).

**Figure 1.**
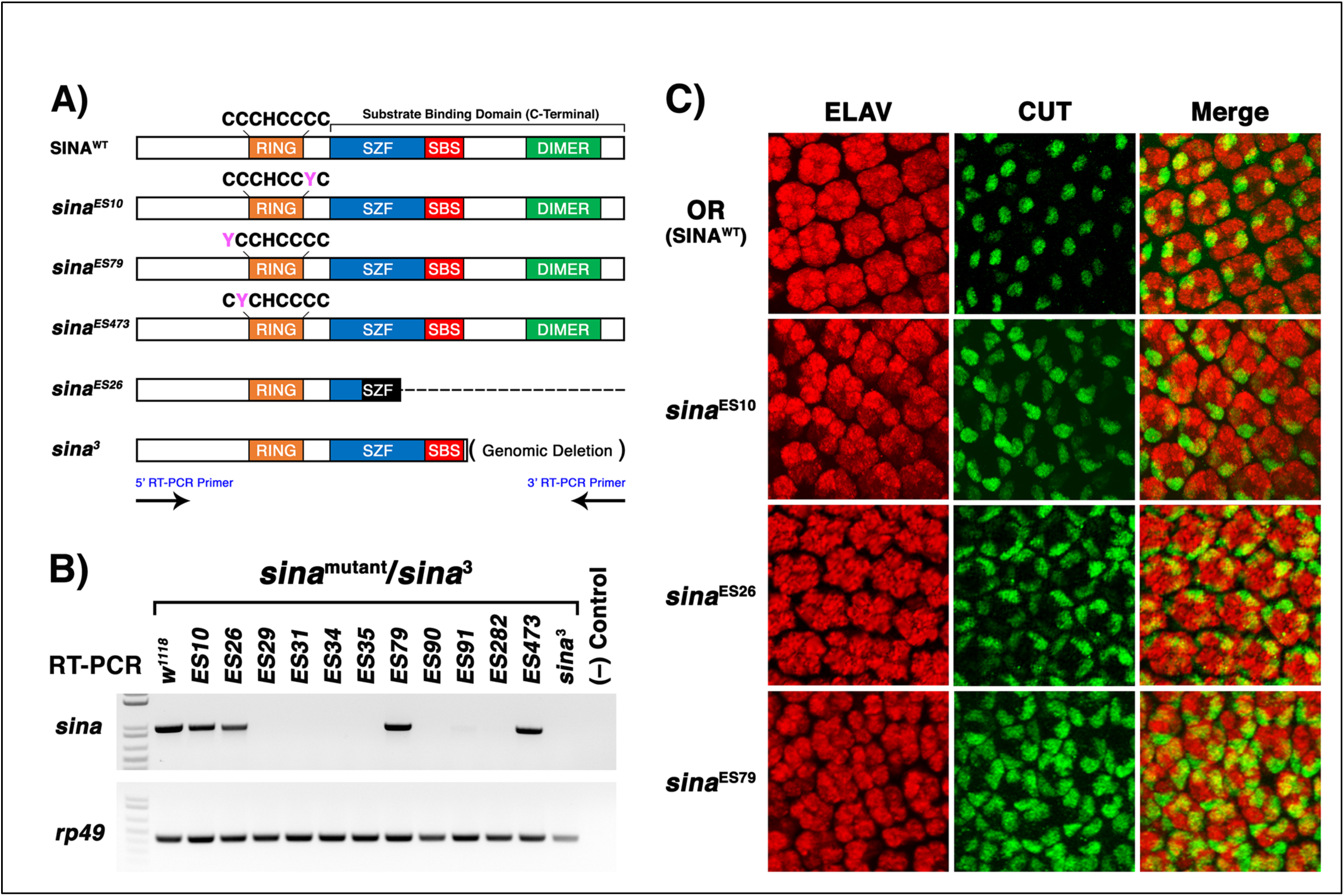
Ethyl methane-sulfonate (EMS) chemical mutagenesis identified the invariable cysteine residues in the RING-domain of SINA whose single point mutation resulted in missing R7 photoreceptor cells and altered ommatidial assembly in *Drosophila* eye development. (**A**) Using ethyl methane-sulfonate (EMS) chemical mutagenesis, 11 newly identified *sina* mutant alleles were isolated from a *GMR-phyl* genetic modifier screen. Among them, three independent single cysteine mutations were identified in the zinc-coordinating RING-domain (*sina*^ES10^, *sina*^ES79^, *sina*^ES473^), while a fourth mutation (*sina*^ES26^) resulted in a 10-bp deletion and a frameshift mutation in the SIAH-type zinc finger (SZF) domain, and thus generated a larger C-terminal truncation than that of the *sina*^3^ mutant allele. (**B**) RT-PCR was performed on the 11 newly identified *sina*^*EMS*^ mutant alleles using the RT-PCR primer pairs at SINA start codon (5’ RT-PCR primer) and stop codon (3’ RT-PCR primer). Four *sina*^*EMS*^ mutant alleles (*sina*^ES10^, *sina*^ES26^, *sina*^ES79^, *sina*^ES473)^ produced *sina* mRNA mutant transcripts in three separate batches of cDNAs synthesized from independent batches of heterogenous *sina*^*EMS*^/*sina*^3^ mutant animals, i.e., 3^rd^ instar wandering mutant long larvae. (**C**) Immunofluorescent staining of ELAV (a neuronal cell marker that marks photoreceptor cells) and CUT (a nonneuronal cell marker that marks cone cells) in developing eye imaginal discs revealed the missing R7, R3 or R4 photoreceptor cells and extra cone cell mutant phenotypes in homozygous *sina*^ES10^ and *sina*^ES79^ mutant animals. Wild type eye discs from the 3^rd^ instar wandering larvae were used as the control.

By using the aforementioned *sina* ORF-specific sequencing analyses, we identified three critical point mutations in the RING domain of SINA from the 3 new *sina*^EMS^ mutant alleles (*sina*^ES10^, *sina*^ES79^, and *sina*^ES473^). The three critical amino acids were at positions of *sina*^C104Y^, *sina*^C73Y^, or *sina*^C76Y^, reflecting a single amino acid substitution in *sina*^ES10^, *sina*^ES79^, or *sina*^ES473^, respectively, as well as a 10 base pair small deletion in *sina*^ES26^, producing a bigger truncation of SINA C-terminal sequence than that of *sina*^3^ mutation (**Figure 1A**). Our phylogenetic analysis showed that each of these cysteine residues (C104Y, C73Y, and/or C76Y) corresponded to an invariable cysteine that is responsible for zinc-coordination in the RING domain of SINA E3 ligase. Our results indicated the critical cysteine residues are required for proper SINA function in relaying RAS signal in *Drosophila* eye development (**Figure 1**) [31, 32].

### The homozygous *sina*^EMS^ mutant larvae, *sina*^ES10^, *sina*^ES26^, or *sina*^ES79^, resulted in loss of photoreceptors, altered cell fates, and disrupted ommatidial assembly in the *Drosophila* developing eye

A majority of 28 newly isolated *sina* mutant alleles are embryonic lethal, resembling that of a large *sina* deletion allele, *sina*^DF(81K72)^ [33]. Using a similar RT-PCR amplification strategy, we designed a series of six paired genomic PCR primers based on the Flybase *sina* genome sequence. These additional primer pairs were designed to be located approximately 500 bp apart in a sequential and stepwise fashion away from the start codon and/or the stop codon of the SINA coding region. An overlapping matrix of RT-PCR synthesis scheme was performed, and we were unable to amplify most of the *sina*^X-ray^ mutant alleles, suggesting that a variety of large genomic deletions in the *sina* gene induced by X-ray mutagenesis did not produce a stable or detectable *sina* mRNA transcript, excluding the sequencing and mutational analyses of these *sina*^X-ray^ mutant alleles. Only three of the 11 new *sina*^EMS^ alleles are homozygous viable as 3^rd^ instar wandering larvae (**Figure 1C**). ELAV is a neuronal cell marker that marks photoreceptor cells in the developing eye imaginal disc, and CUT is a marker for the nonneuronal cone cells in the developing eye. Here, we focused on studying the RAS-dependent neuronal cell fate determination in the developing eyes dissected from *sina*^ES10^, *sina*^ES26^, and *sina*^ES79^ homozygous mutant 3^rd^ instar larvae. Immunofluorescent staining of ELAV and CUT showed that a single point mutation in the RING-domain (*sina*^C104Y^ or *sina*^C73Y^), or a larger C-terminal deletion allele, *sina*^ES26^, impeded R7 photoreceptor cell fate determination in the *Drosophila* eye development. OR (wildtype) larvae were used as the controls to characterize the *sina*^ES10^, *sina*^ES26^, and *sina*^ES79^ mutant phenotypes (**Figures 1A and 1C**).

In the homozygous *sina*^ES10^, *sina*^ES26^, and *sina*^ES79^ larvae, we observed a decrease in the number of ELAV-positive photoreceptor cells and a corresponding increase in the number of CUT-positive cone cells in mutant eye discs (**Figure 1C**). These results indicated that *sina* point mutations resulted in a loss of R7 photoreceptor phenotype and corresponding increased cone cells in the mutant eye discs, suggesting a corresponding cell fate changes due to a loss of SINA function in a single point mutation (*sina*^ES10^ = *sina*^C104Y^, or *sina*^ES79^ = *sina*^C73Y^) (**Figure 1C**). By examining the mutant eye phenotypes of these 3 new *sina* mutant alleles *(sina*^ES10^, *sina*^ES26^, *sina*^ES79^), we found that *sina*^ES79^ point mutation exhibited the strongest mutant phenotype of missing three photoreceptors, R7, R6 and R1, in photoreceptor cell fate determination, while *sina*^ES26^ C-terminal deletion allele exhibited a similar mutant phenotype like that of *sina*^3^ C-terminal deletion allele in *Drosophila* eye development (**Figure 1C** and **Supplemental Figure S2**). Although *sina*^ES473^/sina^3^ heterozygous third instar wandering larvae were viable for RT-PCR analysis, *sina*^ES473^ homozygous larvae were not viable at third instar wandering stages, precluding us from studying its mutant phenotype in the eye imaginal discs.

These results suggested that the invariable cysteine in the zinc-coordinating RING domain is of critical importance to maintain SINA function in relaying active RAS signal in R7 cell fate determination. Comparison of the mutant lethality suggests that the two large-truncated *sina*^2^ and *sina*^3^ mutant alleles may retain partial function since these mutant animals survive until adult stage, despite expressing a C-terminal-deleted SINA protein lacking the dimerization domain. Based on the homozygous larval lethality of these new *sina* mutant alleles *(sina*^ES10^, *sina*^ES79^, or *sina*^ES473^), we concluded that without any of the three critically important cysteine residues in its RING-domain, SINA cannot function properly, suggesting that C104Y, C73Y, or C76Y is indispensable for proper SINA function (**Figure 1**). The classic and complementary cell fate changes observed in these *sina* cysteine point mutants suggested that proper zinc-coordination in the RING-domain is critical for SINA’s gatekeeper function in relaying RAS-mediated signal transduction in R7 photoceptor cell fate determination (**Figure 1C**) [25, 33, 34].

### *Drosophila* SINA and human SIAH1/2 are functionally conserved in *Drosophila* eye development

Supported by a high degree of amino acid identity and cross-species protein sequence conservation in metazoan SINA/SIAH proteins, we hypothesized that human SIAHs are likely to play a similar gatekeeper function as *Drosophila* SINA by serving as the key downstream signaling hub essential for active K-RAS signal transduction [28-30, 32]. To test this hypothesis, we examined the functional conservation of *Drosophila* SINA and human SIAH1/2 *in vivo*.

Importantly, we conducted a cross-species genetic analysis to determine whether targeted expression of SIAH^PD^, a known SIAH inhibitor, with potent anti-K-RAS and anticancer efficacy in xenograft mouse models of human pancreatic and lung cancer [29, 30], could also inhibit the RAS-dependent neuronal cell fate determination in *Drosophila.*

These bioengineered SINA^PD^/SIAH^PD^ mutant proteins feature two Cys to Ser point mutations in the invariable cysteine (Cys) residues of the zinc-coordinating RING domain of *Drosophila* SINA (SINA^C73S-C76S^) and human SIAH1/2 protein (SIAH1^C41S-C44S^ and SIAH2^C80S-C83S^). These potent SIAH inhibitors effectively inactivate its E3 ligase activity and block human SIAH1/2 self-ubiquitination and substrate degradation, creating a proteolysis-deficient (PD) full-length SIAH1/2 mutant protein [29, 30]. The SIAH1/2 inhibitors can bind to their respective substrates, but cannot degrade them, effectively ablating SIAH-mediated substrate proteolysis [29]. We showed that the RING-domain point mutants, SIAH1^PD^/SIAH2^PD^, were highly effective to eradicate and abolish oncogenic K-RAS-driven tumorigenesis in xenograft mouse models [29, 30]. Here, we aimed to determine whether these bioengineered SINA/SIAH inhibitors might be able to mimic the key aspects of *sina*^loss of function^ mutant phenotype by blocking endogenous SINA function and impeding RAS-dependent photoreceptor and PNS development in *Drosophila.*

To compare the mutant phenotypes of *sina*^EMS^ point mutant alleles to those of the bioengineered SINA^PD^/SIAH^PD^ constructs, we established a panel of transgenic lines that express either the wild type (WT) or proteolysis deficient (PD) point mutations of *Drosophila* SINA and the human SINA homologs SIAH1 and SIAH2. We first cloned, sequence verified, injected, and generated a complete panel of *UAS-sina*^WT^/*UAS-sina*^PD^ and *UAS-SIAH*^WT^/*UAS-SIAH*^PD^ transgenic flies. We then examined the perturbation of RAS signaling in photoreceptor and PNS development in response to ectopic expression of *Drosophila* SINA^WT^/human SIAH1/2^WT^, as well as the cellular defects due to expression of the SINA/SIAH inhibitors, *Drosophila* SINA^PD^/human SIAH1/2^PD^, under multiple tissue-specific GAL4-drivers *in vivo* (**Figures 2, 3, 4, 7, 8** and **9**).

**Figure 2.**
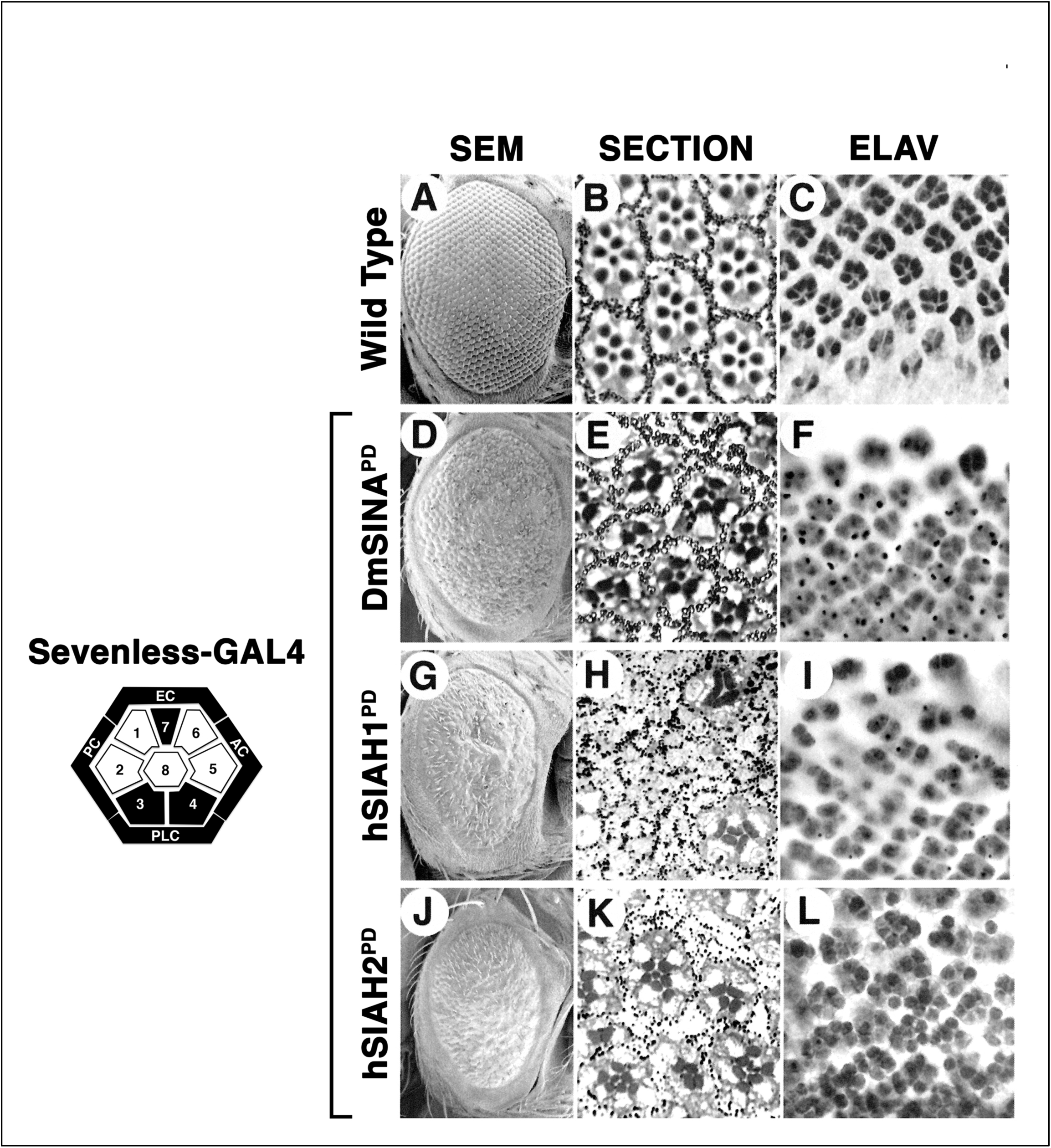
Ectopic expression of *Drosophila* SINA^PD^ (Dm SINA^PD^), human SIAH1^PD^ (hSIAH1^PD^) and human SIAH2^PD^ (hSIAH1^PD^) under the control of *Sevenless-GAL4* blocked photoreceptor cell differentiation, and resulted in altered ommatidial assembly and rough eye phenotypes. The *Sevenless-GAL4* expression pattern is illustrated by a schematic diagram of a developing ommatidia. The R7, R3, R4 photoreceptors and four cone cells that are specifically targeted by this eye-specific driver, *Sevenless-GAL4*, are highlighted in black color. The scanning electron microscopy (SEM) (**A**), adult retina section (**B**), and ELAV staining of the photoreceptor cell clusters (**C**) in wildtype flies are shown as the normal controls. Ectopic expression of the proteolysis-deficient (PD) DmSINA^PD^, hSIAH1^PD^, and hSIAH2^PD^ under the control of *Sevenless*-*GAL4* resulted in rough eyes, and loss of R7, R3 and R4 photoreceptor phenotypes, and disorganization of ommatidial assembly as shown by scanning electron microscopy (SEM) (**D, G** and **J**) and sectioning of the adult retina (**E, H** and **K**). Immunohistochemical staining of ELAV, a neuronal cell marker that marks the photoreceptor cells in the developing eye imaginal discs, indicates the loss of R7, R3 and R4 photoreceptor phenotypes (**F**), increased photoreceptor cell death as marked by the dense dots and puncta, and severe defects in photoreceptor cell differentiation and photoreceptor cluster assembly (**I** and **J**) in response to the SINA blockade due to SINA^PD^, SIAH1^PD^, and SIAH2^PD^ ectopic expression in a subset of photoreceptor cells (R7, R3, and R4) in the developing eyes.

**Figure 3.**
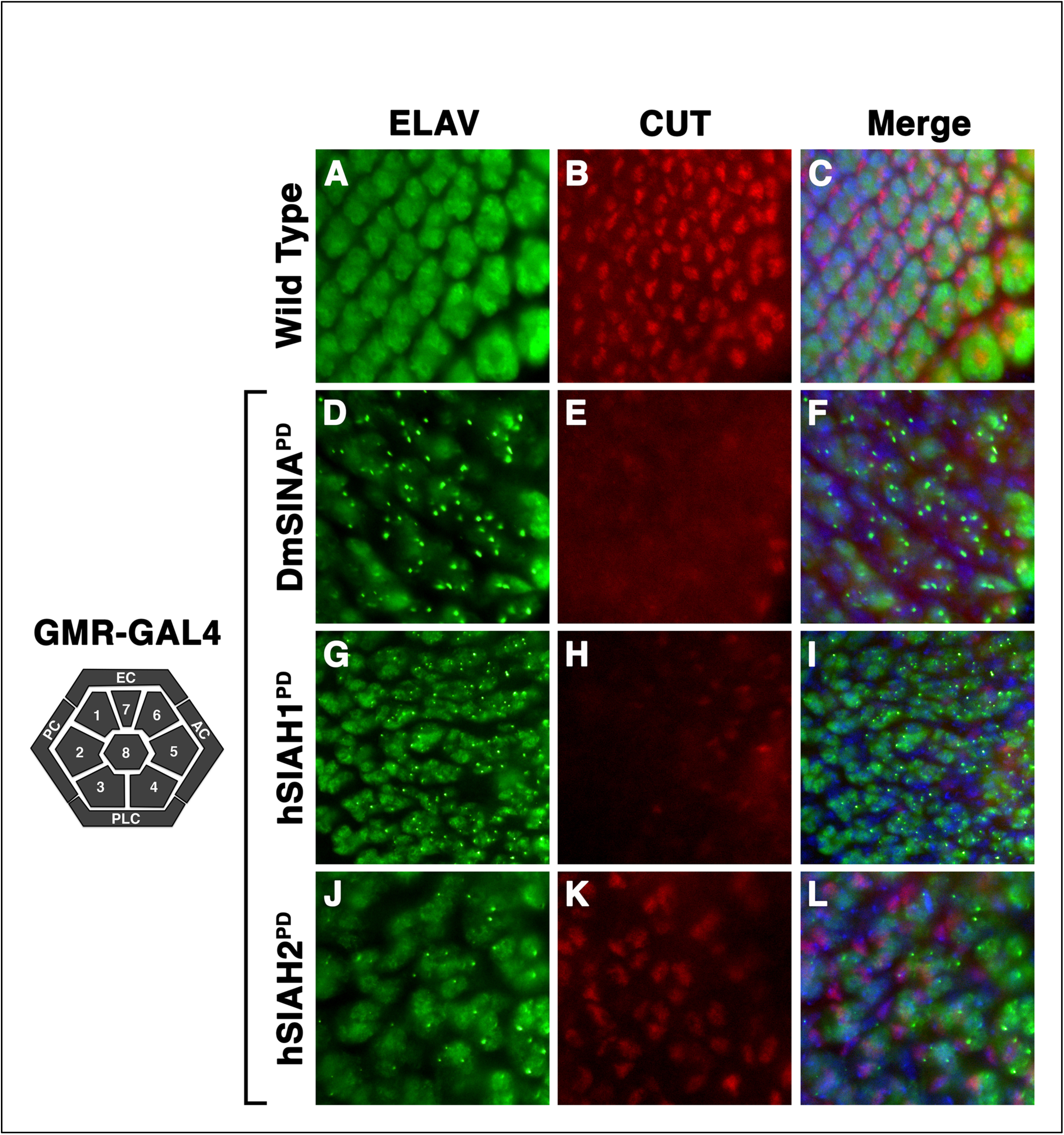
Ectopic expression of *Drosophila* SINA^PD^ (Dm SINA^PD^), human SIAH1^PD^ (hSIAH1^PD^) and human SIAH2^PD^ (hSIAH1^PD^) under the control of *GMR-GAL4* blocked photoreceptor cell differentiation, and resulted in altered ommatidial assembly and rough eye phenotypes. The *Glass Multimer Reporter (GMR)*-*GAL4* expression pattern is illustrated by a schematic diagram of a developing ommatidia. All photoreceptors and cone cells that are specifically targeted by this eye-specific driver, *GMR-GAL4*, are highlighted in dark gray color. ELAV is a neuronal cell marker that marks photoreceptor cells of the developing eye imaginal disc (Green), CUT is a marker for the nonneuronal cone cells of the developing eye (red), and Hoechst 33342 is a DNA dye that marks nuclei (blue). Immunofluorescent staining of ELAV (**A**) and CUT (**B**) and montage (**C**) in the wildtype developing eye imaginal discs of third instar wondering larvae are shown as the controls. Ectopic expression of the proteolysis-deficient (PD) DmSINA^PD^, hSIAH1^PD^, and hSIAH2^PD^ under the control of *GMR*-*GAL4* resulted in rough eyes, loss of photoreceptor and cone cell phenotypes, increased cell death as marked by the dense dots and puncta, and disorganization of ommatidial assembly. Immunofluorescent staining of reduced ELAV (**D, G** and **J**) and reduced CUT (**E, F** and **K**) resulted from ectopic expression of the SINA inhibitors, SINA^PD^ (**D, E** and **F**), SIAH1^PD^ (**G, H** and **I**), and SIAH2^PD^ (**J, K** and **L**) under the control of *GMR-GAL4* in all cells in the developing eyes are shown.

**Figure 4.**
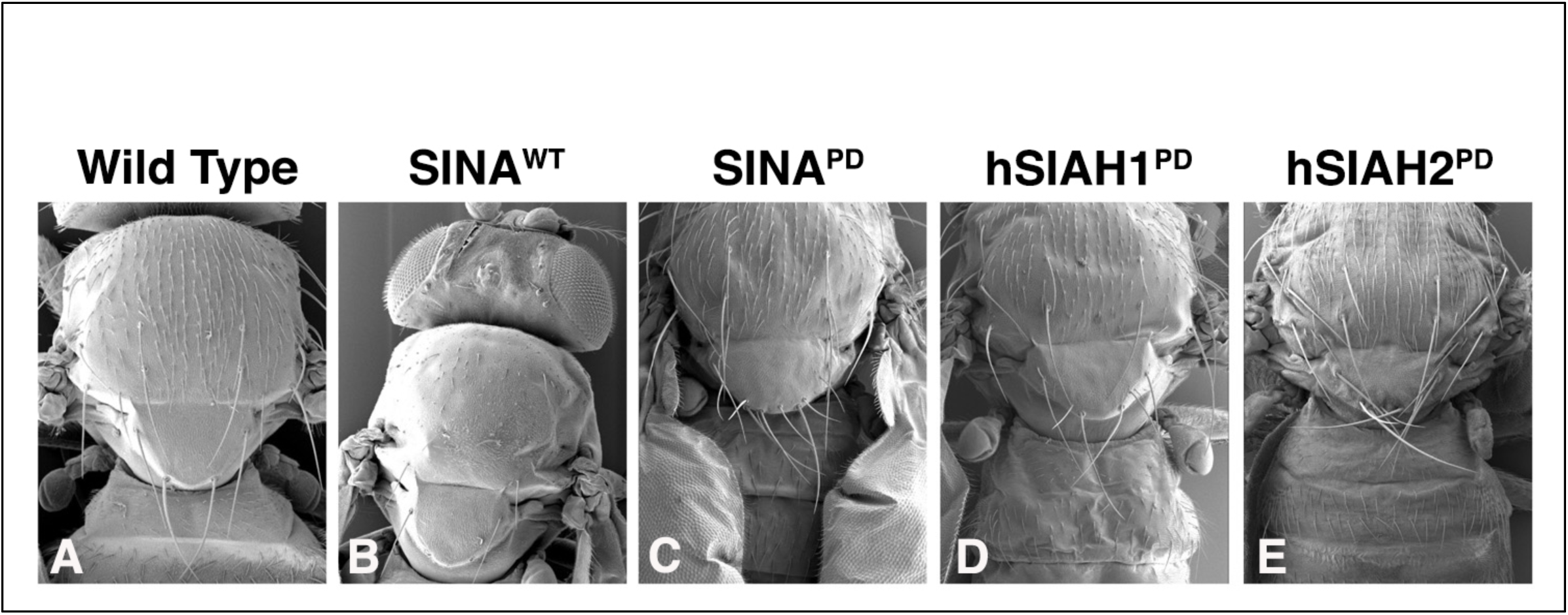
SINA plays an important role in mechanosensory neuronal development in *Drosophila*. The scanning electron microscopy (SEM) of macrochaetes (mechanosensory bristles on the fly notum) in wildtype flies is shown (**A**). Ectopic expression of SINA^WT^ under the control of *Dpp-GAL*4 driver results in missing macrochaetes and almost bald notum phenotype (**B**). In contrast, ectopic expression of the proteolysis-deficient (PD) *Drosophila* SINA^PD^ (**C**), human SIAH1^PD^ (**D**) and human SIAH2^PD^ (**E**) under the control of the *Dpp-GAL*4 driver all result in supernumerary macrochaete formation and “hairy” notum phenotypes.

**Figure 5.**
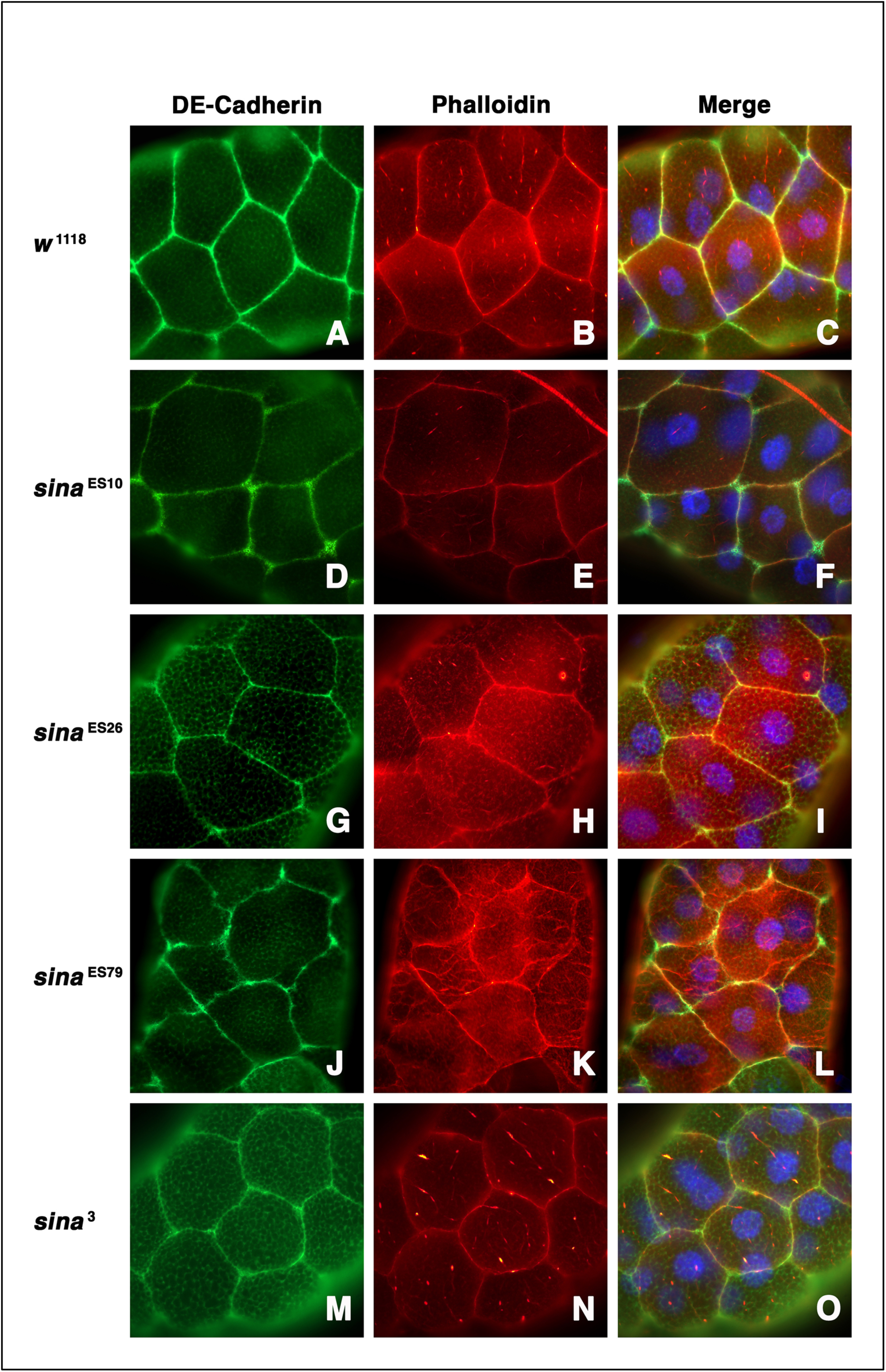
*sina* mutant epithelial cells exhibited defective cell junctions, abnormal and discontinuous DE-Cadherin staining, and rounded-up cell morphology in the larval salivary glands. Salivary glands dissected from the third instar wandering larvae were stained with DE-Cadherin (Green), Phallodin (F-actin, Red), and Hoechst (DNA/nuclei, Blue). The cell junction protein, DE-Cadherin (**A**), actin cytoskeleton (**B**), and the merged image (**C**), that outlined the plasma membrane of adjacent epithelial cells in the wildtype larval salivary gland, are shown as the controls. Immunofluorescent staining of *sina*^ES10^, *sina*^ES26^, *sina*^ES79^ and *sina*^3^ mutant epithelial cells showed defective cell junction and disconnected DE-Cadherin staining (**D, G, J**, and **M**), defective F-actin cytoskeleton network (**E, H, K**, and **N**), defective cell junction and abnormal cell shape as shown in the merged images (**F, I, L**, and **O**) in these *sina* mutant larval salivary glands. Notedly, *sina*^ES79^ has the strongest mutant phenotypes among the 4 *sina* homozygous mutant animals as shown in here.

**Figure 6.**
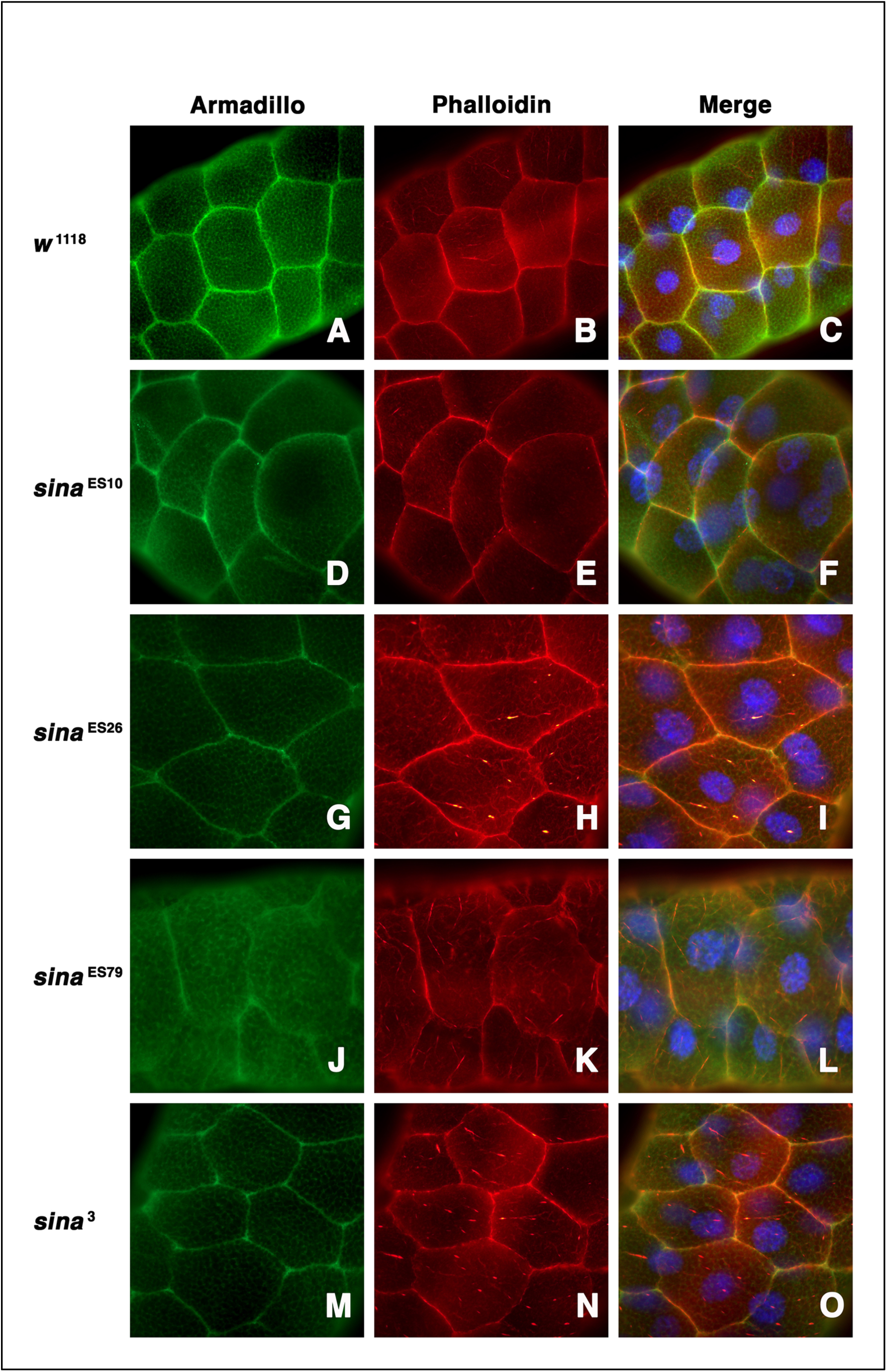
*sina* mutant epithelial cells exhibited defective cell shape, abnormal and mis-localized Armadillo staining, and disorganized actin cytoskeleton in the larval salivary glands. Salivary glands dissected from the third instar wandering larvae were stained with Armadillo (Gree), Phalloidin (F-actin, Red), and Hoechst (DNA/nuclei, Blue). The cell junction protein, Armadillo (**A**), actin cytoskeleton (**B**), and the merged image (**C**), that outlined the plasma membrane of adjacent epithelial cells in the wildtype larval salivary gland, are shown as the controls. Immunofluorescent staining of *sina*^ES10^, *sina*^ES26^, *sina*^ES79^ and *sina*^3^ mutant epithelial cells showed defective, mis-localized, and disintegrated Armadillo staining (**D, G, J**, and **M**), disorganized F-actin cytoskeleton network (**E, H, K**, and **N**), defective cell junction and rounded-up cell morphology as shown in the merged images (**F, I, L**, and **O**) in these *sina* mutant larval salivary glands. Notedly, *sina*^ES79^ has the strongest mutant phenotypes among the 4 *sina* homozygous mutant animals as shown in here.

**Figure 7.**
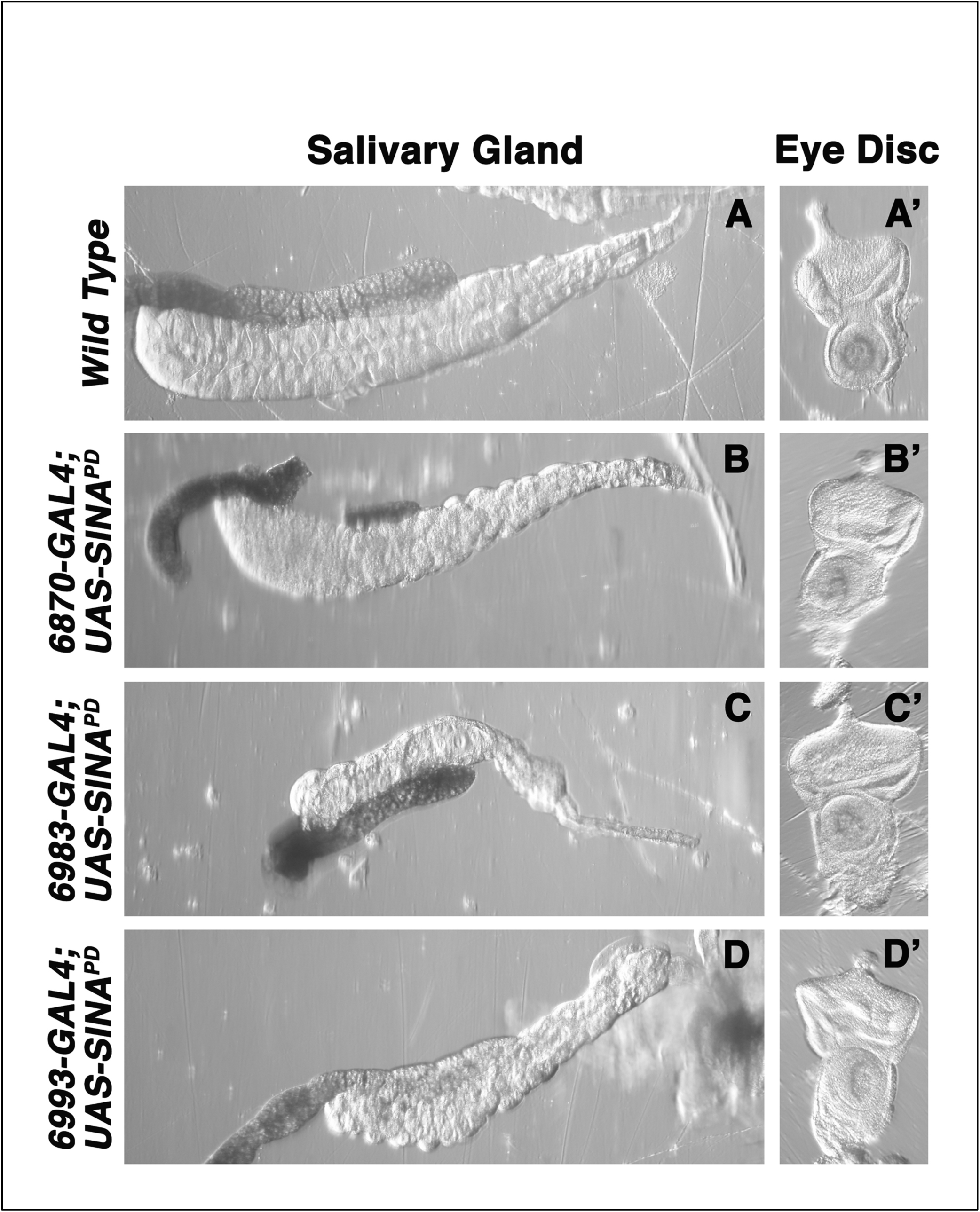
Ectopic expression of SINA inhibitor, SINA^PD^, under the control of three salivary gland-specific GAL4 drivers resulted in cellular defects, junctional disorganization, or tissue atrophy in the larval salivary gland. The normal epithelial cells display highly organized cell junction and hexagonal epithelial cell morphology in the wildtype type larval salivary glands under a dissection microscope (**A**). Ectopic expression of SINA^PD^ under the control of three salivary gland (SG)-specific GAL4-drivers resulted in cellular defects, junctional disorganization, or tissue atrophy in the larval salivary gland. *6870-GAL4* is a weak SG-driver, *6983-GAL4* is a strong SG-driver, and *6993-GAL4* is an intermediate SG-driver. Ectopic expression of SINA^PD^ under the control of the weak *6870-GAL4* resulted in cellular defects, salivary gland disorganization, and mild tissue atrophy (**B**). Ectopic expression of SINA^PD^ under the control of the strong *6983-GAL4* resulted in small, disformed, and dying salivary gland with severe tissue atrophy (**C**). Ectopic expression of SINA^PD^ under the control of the intermediate *6993-GAL4* resulted in defective, disorganized, and atrophied salivary gland (**D**). Eye discs dissected from the same transgenic animals were used as the internal controls to show the precise timing of the identical developmental stages from which the mutant salivary glands were dissected to examine the tissue morphology and IF staining from the same transgenic animals (**A’, B’, C’** and **D’**).

**Figure 8:**
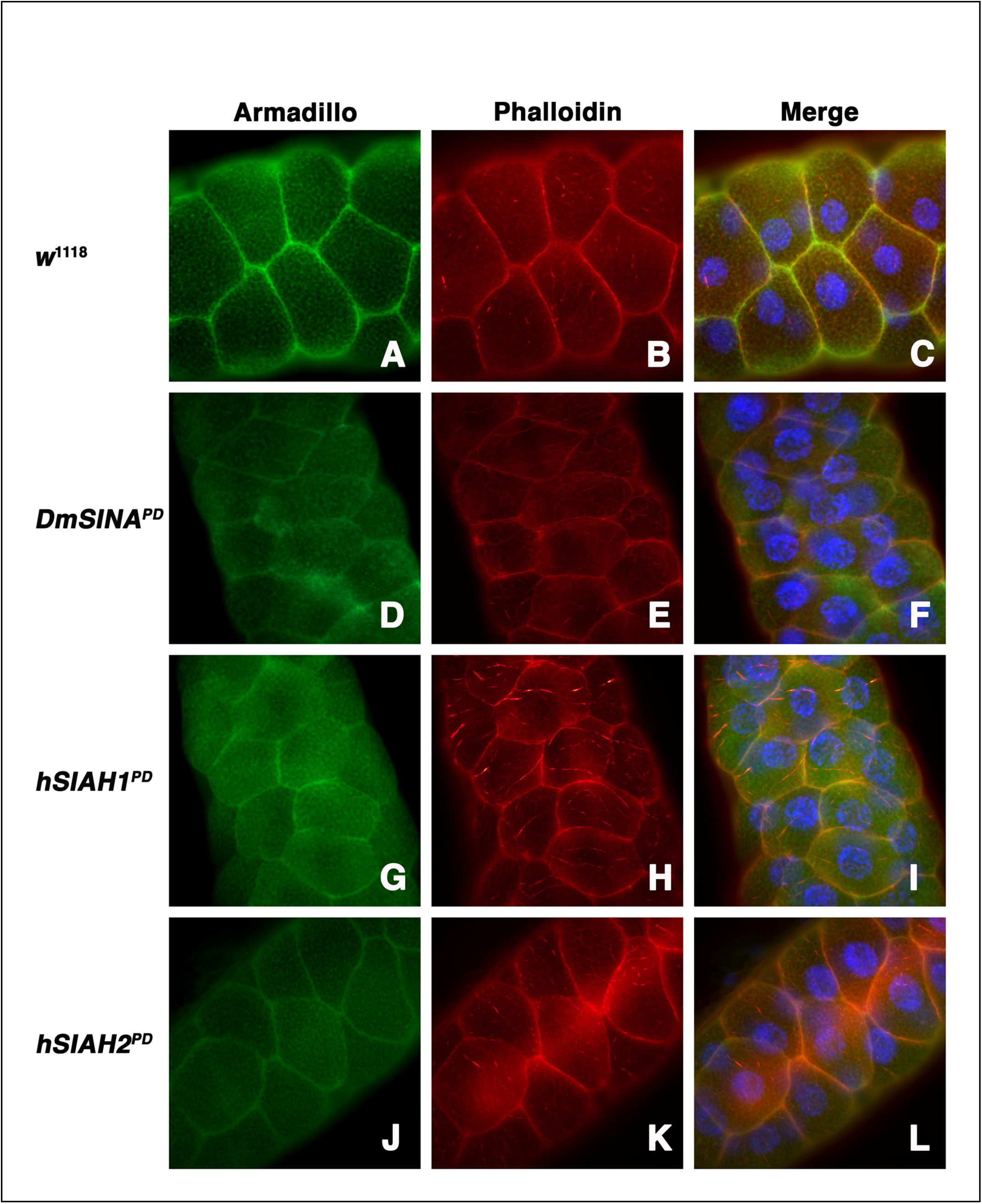
Ectopic expression of the SINA/SIAH inhibitors (DmSINA^PD^, hSIAH1^PD^, and hSIAH2^PD^), under the control of a SG-specific *6870-GAL4*, resulted in defective cell junction, miss-localized Armadillo staining, and abnormal cell shape in SINA-deficient epithelial cells in the transgenic salivary glands. Salivary glands dissected from the third instar wandering larvae were stained with Armadillo (Green), Phalloidin (F-actin, Red), and Hoechst (DNA/nuclei, Blue). Immunofluorescent staining of the cell junction protein, Armadillo (**A**), F-actin (**B**), and the merged image (**C**), in the wildtype larval salivary gland, are shown. Immunofluorescent staining of transgenic salivary glands, resulted from ectopic expression of DmSINA^PD^, hSIAH1^PD^ and hSIAH1^PD^ under a weak SG-driver, *6870-GAL4*, revealed defective cell junction, mislocalized Armadillo staining (**D, G**, and **J**), altered cell shape and distorted actin cytoskeleton network (**E, H**, and **K**), in the SINA-deficient epithelial cells (**F, I**, and **L**).

**Figure 9:**
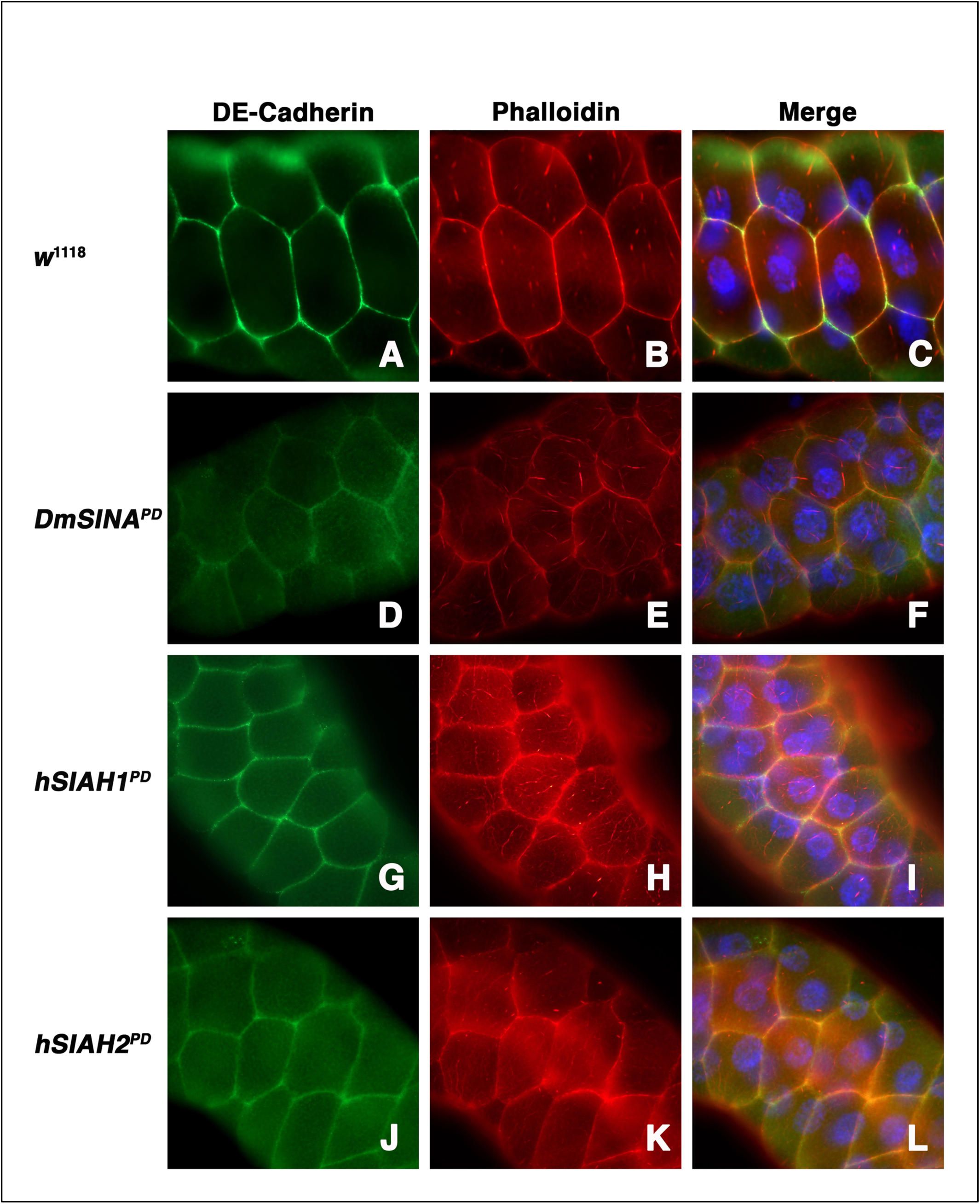
Ectopic expression of the SINA/SIAH inhibitors (DmSINA^PD^, hSIAH1^PD^, and hSIAH2^PD^), under the control of a SG-specific *6870-GAL4*, resulted in defective cell junction, reduced DE-Cadherin staining, and abnormal cell shape in SINA-deficient epithelial cells in the transgenic salivary glands. Salivary glands dissected from the third instar wandering larvae were stained with DE-cadherin (Green), Phalloidin (F-actin, Red), and Hoechst (DNA/nuclei, Blue). Immunofluorescent staining of the cell junction protein, DE-cadherin (**A**), F-actin (**B**), and the merged image (**C**), in the wildtype larval salivary gland, are shown. Immunofluorescent staining of transgenic salivary glands, resulted from ectopic expression of DmSINA^PD^, hSIAH1^PD^ and hSIAH1^PD^ under a weak SG-driver, *6870-GAL4*, revealed defective cell junction, abnormal DE-cadherin staining (**D, G**, and **J**), altered cell shape and distorted actin cytoskeleton network (**E, H**, and **K**), in the SINA-deficient epithelial cells (**F, I**, and **L**).

### Ectopic expression of *Drosophila* SINA^PD^ and human SIAH1^PD^/SIAH2^PD^, under the control of eye-specific GAL4 promoters disrupted photoreceptor cell fate determination

The normal composition of the *Drosophila* photoreceptors and cone cells in the wildtype adult and developing eye is shown (**Figures 2A, 2B, 2C, 3A, 3B**, and **3C**). Ectopic expression of SINA^WT^/SIAH^WT^ under eye-specific *GMR-* or *Sev-GAL4* promoter resulted in mild or medium rough eye phenotype, without altering photoceptor cell fate determination in the developing eyes as indicated by ELAV or CUT immunofluorescent (IF) staining (data not shown). These results are consistent with the role of SINA as a critical permissive factor of RAS signaling transmission, but not a driver of RAS-dependent neuronal cell differentiation in the eye development *in vivo* [24, 25, 33, 41]. Here we focused on characterizing the strong rough eye phenotypes of expressing the putative SINA/SIAH inhibitors, i.e., SINA^PD^/SIAH1^PD^/SIAH2^PD^, in *Drosophila* eye development *in vivo*.

Firstly, targeted expression of SINA^PD^ inhibitor to a subset of photoreceptor cells, R7, R3 and R4, using the *Sevenless* (*Sev*)*-GAL4* driver resulted in a loss of these three photoreceptor cells and a strong rough eye phenotype in adult flies (**Figure 2D**). Sectioning of the rough eyes revealed a disorganization of the highly structured ommatidial array as well as three missing photoreceptor cell mutant phenotype (**Figure 2E**). Consistent with the adult eye phenotypes, immunofluorescence (IF) staining of larval eye imaginal discs revealed a loss of the targeted R7, R3 and R4 photoreceptor cells in the developing eyes of the transgenic animals (**Figure 2F**). The three SINA-deficient photoreceptors were dead as marked by the three small, dense punctae in each ommatidial cluster in the developing eye (**Figure 2F**). Similarly, ectopic expression of human SIAH1^PD^ or human SIAH2^PD^ under the control of *Sev-GAL4* driver resulted in strong rough eye phenotypes and a loss of multiple photoreceptor mutant phenotype in adult flies (**Figures 2G and 2J**). Since proper SINA function is dependent on SINA dimerization, it is conceivable that these SINA inhibitors, SINA^PD^/SIAH^PD^, bind to endogenous SINA to shut down its biological activities required for photoreceptor cell fate determination in the *Drosophila* eye development. Sections of adult eyes expressing human SIAH1^PD^ or human SIAH2^PD^ resulted in a highly disorganized ommatidial array, and missing photoreceptor cells in the adult eye (**Figures 2H and 2K**). Immunofluorescent staining of eye imaginal discs confirmed the missing photoreceptor mutant phenotypes and highly disorganized ommatidia in the developing eyes of these transgenic animals expressing hSIAH1^PD^/hSIAH2^PD^ (**Figures 2I and 2L**). Based on their extensive phylogenetic and evolutionary conservation, and the similar mutant phenotypes resulted from expressing SINA^PD^/SIAH^PD^ in a subset of photoreceptor cells, it is likely that human SIAHs and *Drosophila* SINA are functionally conserved to impede endogenous *Drosophila* SINA activity to successfully block proper RAS signal transduction critical for R7, R3 and R4 neuronal cell differentiation *in vivo*.

Secondly, the Glass Multimer Reporter (*GMR*)-*GAL4* enhancer drives expression of exogeneous UAS-gene to all photoreceptor cells and surrounding cone cells in the developing eye, though *GMR-GAL4* expression is to a lesser degree compared with that of *Sev-GAL4* [42, 43]. GMR-driven expression of SINA^PD^, SIAH1^PD^, and SIAH2^PD^ in the eye resulted in strong rough eye phenotypes in adult flies. Dissection and immunofluorescent staining of transgenic eye discs of the resultant wandering 3^rd^ instar larvae revealed blocked photoreceptor development, reduced numbers of photoreceptors and cone cells, increased cell death, and disorganized ommatidial assembly upon SINA blockade (**Figure 3**). In the wild type eye discs, ELAV and CUT staining shows a stereotypical eight photoreceptor assembly surrounded by four cone cells in a highly ordered and organized fashion in the developing eye (**Figures 3A, 3B** and **3C**). GMR-driven expression of SINA^PD^ (**Figures 3D, 3E** and **3F**) and SIAH1^PD^ (**Figures 3G, 3H** and **3I**) resulted in disorganization of the ommatidial structure, reduced ELAV-positive photoreceptor cells, featuring a punctate like ELAV staining pattern indicative of photoreceptor cell death, and loss of the CUT-positive cone cells. GMR-driven expression of SIAH2^PD^ also resulted in disorganization of the ommatidial structure with fewer punctae in the ELAV staining and fewer ELAV and CUT cells in each ommatidium (**Figure 3J, 3K** and **3L**). These results indicated that ectopic expression of these SINA/SIAH inhibitors, *Drosophila* SINA^PD^/human SIAH^PD^, blocked photoreceptor and cone cell development, induced neuronal cell death, mimicking aspects of strong *sina*^loss of function^ mutant phenotypes in RAS-dependent photoreceptor cell differentiation *in vivo* (**Figures 1, 2 and 3**).

### Ectopic expression of *Drosophila* SINA^WT/PD^, human SIAH1^WT/PD^ and SIAH2^WT/PD^ under *Dpp-GAL4* altered mechanosensory bristle development *in vivo*

The *sina*^Loss of function^ mutant phenotypes are not limited to R7 photoreceptor cells. In hypomorphic *sina* mutant flies, the lack of macrochaete and microchaete and missing sensory neurons were another prominent feature of the *sina*^2^ or *sina*^3^ homozygous mutant phenotype, suggesting a role of SINA in *Drosophila* peripheral nervous system (PNS) development [33]. Macrochaete are mechanosensory bristles derived from epidermal cells and coupled with sensory neurons [44-46]. For mechanosensory bristle development, sensory organ precursor (SOP) undergoes two rounds of asymmetric cell divisions and confers four distinct cell fates to the daughter cells [47-52]. The *dpp*^*40C6*^*-GAL4* enhancer drives gene expression to the wing imaginal disc, resulting in wing and notum phenotypes [53]. The wildtype macrochaetes and microchaetes on a normal notum was shown (**Figure 4A**). Ectopic expression of SINA^WT^ under the control of *dpp-GAL4* driver resulted in a dramatic loss of macrochaetes and microchaetes on the notum (**Figure 4B**). In contrast, ectopic expression of DmSINA^PD^/hSIAH1/2^PD^ under the control of *dpp-GAL4* driver resulted in the formation of extranumerary macrochaetes (**Figures 4C, 4D and 4E**). Like its *Drosophila* counterparts, ectopic expression *SINA*^WT/PD^/*SIAH*^WT/PD^ under *Dpp-GAL4* driver resulted in opposing phenotypes, i.e., the missing (WT) and extra (PD) sensory bristles (e.g. macrochaetes), on the notum of the transgenic flies, suggesting that SINA^PD^/SIAH^PD^ expression alters the PNS neuronal cell fate determination in *Drosophila* notum (**Figure 4**). Based on the similar macrochaete phenotypes observed in the cross-species analyses, it is likely that that *Drosophila* SINA and human SIAH1/SIAH2 may be functionally conserved in *Drosophila* PNS development.

### *The sina* mutant epithelial cells displayed defects in cell shape, cytoskeleton network, and cell junction in the larval salivary glands

We reported that SIAH-deficiency altered focal adhesion, cell junction, cell shape and cell attachment in human cancer cells (Tang manuscript in revision). As a single layer of epithelial cells, the *Drosophila* salivary gland provides a unique opportunity to study the role of SINA on epithelial cell biology *in vivo.* In wild type animals, Armadillo (*Drosophila* β-catenin homolog) and DE-Cadherin are two cell junction proteins whose membrane-specific expression highlights and demarcates the borders of adjacent epithelial cells in the larval salivary glands (**Figures 5A, 5B** and **5C, 6A, 6B** and **6C**). To examine the effect of *sina*^loss of function^ mutant phenotypes *in vivo*, we conducted IF staining of DE-cadherin and Armadillo, and actin cytoskeleton network in *sina* mutant epithelial cells of the larval salivary glands freshly dissected from wandering 3^rd^ instar larvae (**Figures 5** and **6**). We found the *sina*^ES10^, *sina*^ES26^, *sina*^ES79^, and *sina*^3^ mutant epithelial cells displayed a rounded-up cell morphology, detached and distorted cell junction, decreased and discontinuous cell attachment, mis-localized and punctated staining of Armadillo and DE-Cadherin in the *sina*^ES10^ (**Figures 5D, 5E, 5F, 6D, 6E**, and **6F**), *sina*^ES26^ (**Figures 5G, 5H, 5I, 6G, 6H**, and **6I**), *sina*^ES79^ (**Figures 5J, 5K, 5L, 6J, 6K**, and **6L**), and *sina*^3^ **(Figures 5M, 5N, 5O, 6M, 6N** and **6O**) mutant salivary glands. Furthermore, the *sina*^ES10^, *sina*^ES26^, *sina*^ES79^, and *sina*^3^ mutant epithelial cells displayed a defective cell border, cell detachments, altered cell shape, weakened cell junction, and disorganized cytoskeletal network (actin and microtubule) when compared to the normal actin and tubulin staining from the regular hexagon-shaped epithelial cells in wildtype salivary glands as the controls (**Figures 5** and **6**). Notedly, the cellular defects in the *sina*^ES79^ mutant epithelial cells (**Figures 5J, 5K, 5L, 6J, 6K**, and **6L**) are much stronger than those of the *sina*^3^ mutant epithelial cells (**Figures 5M, 5N, 5O, 6M, 6N** and **6O**).

### Ectopic expression of the SINA/SIAH inhibitors in the larval salivary gland disrupted epithelial cell junctions, altered cell shape, and induced cell death and tissue atrophy

We next examined whether SINA inhibition due to ectopic expression of the SINA/SIAH inhibitor would mimic aspects of *sina*^loss of function^ mutant phenotypes on epithelial cell biology. To block endogenous SINA function in the epithelial cells of larval salivary glands, we expressed *Drosophila* SINA^PD^, human SIAH1^PD^ and SIAH2^PD^, under the control of three distinct salivary gland (SG)-specific GAL4 drivers that express the transgene at different levels to examine the effect of SINA-blockade on salivary gland development (**Figures 7, 8** and **9**). Low-level expression of SINA^PD^ under a weak SG-specific driver, *6870-GAL4*, resulted in a mild SG mutant phenotype (**Figure 7B**). High-level expression of SINA^PD^ under strong SG-specific drivers, *6983-GAL4* and *6993-GAL4*, resulted in strong SG mutant phenotypes with highly atrophied salivary glands (**Figures 7C** and **7D**). It is clear that high-level SINA^PD^ expression induced massive cell death and severe tissue atrophy in the salivary glands of transgenic animals (**Figure 7**). The stereotypic ommatidial pattern formation of the *Drosophila* eye development were included as internal age controls to closely match the identical timing of SG development from these transgenic animals of the same age (**Figures 7A’, 7B’, 7C’** and **7D’**). Hence, the 3^rd^ instar eye imaginal discs with the comparable rows of photoreceptor clusters dissected from the different transgenic animals were used to show the precise timing of the identical developmental stages from which the mutant salivary glands were dissected to show the tissue-specific SG destruction and organ atrophy is GAL4-specific, SINA^PD^/SIAH^PD^-dependent, and non-cell autonomous. Similar cellular phenotypes and tissue atrophies were observed when the human SIAH inhibitors, SIAH1^PD^ and SIAH2^PD^, were expressed under the control of these SG-specific drivers (*6870-GAL4, 6983-GAL4*, and *6993-GAL4*) as well (**Supplemental Figures S3** and **S4**). These results suggested that epithelial cell patterning of the salivary gland is a good cell-based system to study SINA-dependent epithelia cell biology *in vivo.*

To minimize studying any cellular artifacts introduced by atrophied and dying salivary gland cells, we focused our efforts on studying the cellular defects of expressing a potent SINA/SIAH inhibitor in the epithelial cells under the control of *6870-GAL4*. This weak SG-specific driver expresses *Drosophila* SINA^PD^, human SIAH1^PD^, or human SIAH2^PD^ at a low-level that does not induce epithelial cell death and tissue atrophy in the transgenic salivary glands of the same age. Again, endogenous Armadillo (*Drosophila* β-catenin homolog) and DE-Cadherin is localized to the plasma membrane that demarcates highly organized epithelial cell junctions in wildtype larval salivary gland (**Figures 8A, 8B, 8C, 9A, 9B**, and **9C**). In contrast, low-level expression of SINA^PD^, hSIAH1^PD^, or hSIAH2^PD^ under *6870-GAL4* altered cell shape, reduced Armadillo/DE-Cadherin membrane localization, and increased Armadillo/DE-Cadherin cytoplasmic mislocalization in the transgenic salivary glands (**Figures 8D, 8G, 8J, 9D, 9G** and **9J**).

Immunofluorescent staining of Phalloidin (F-actin) revealed defective cell junctions, cell borders, actin cytoskeleton, and abnormal cell shape and morphology in response to SINA blockade, i.e., SINA^PD^/SIAH^PD^ inhibitor expression in the salivary gland epithelial cells (**Figures 8E, 8F, 8H, 8I, 8K, 8L, 9E, 9F, 9H, 9I, 9K** and **9L**).

## Discussion

Underscoring the central importance of SINA biology in the RAS pathway is its extraordinary degree of evolutionary conservation from invertebrates to vertebrate species, with over 83% amino acid identity shared between *Drosophila* SINA and human SINA homologs (SIAHs) [29, 31, 40]. Our newly identified large *sina* mutant collection is particularly valuable and medically relevant for the scientific community, pertinent to controlling undruggable oncogenic RAS hyperactivation in cancer biology and oncology. The complementation studies suggested that the previously published five *sina* mutants (*sina*^1^, *sina*^2^, *sina*^3^, *sina*^4^, and *sina*^5^) are all weak hypomorphic alleles, i.e., partial loss-of-function alleles, and *sina*^*3*^ and *sina*^*2*^ are not the null alleles as previously suggested [33]. With the new revelation that the *sina* C-terminal truncated alleles are actually hypomorphic alleles, there is a need to re-evaluate the current anti-SIAH strategy used in the mammalian field. **Firstly**, the *Siah2*^*-/-*^ knockout mice were generated using a similar C-terminal protein truncation strategy to mutate the *Siah2* gene to generate a “null” allele [54]. However, our SINA mutagenesis study indicated that C-terminal SIAH2 truncation will not generate a null allele, i.e., the complete loss of function mutation, in mice. Hence, homozygous *Siah2*-knockout mutant mice with a C-terminal truncation at SIAH2 amino acid residue 180, mimicking *Drosophila* SINA^2^ or SINA^3^ mutant protein with the comparable C-terminal truncations, is likely a hypomorphic allele. **Secondly**, X-ray structural and biochemical analysis of mouse SIAH1a and human SIAH1 utilized the N-terminal truncated forms of SIAH1, missing ∼80-90 amino acids to delete the entire catalytic RING-domain (missing nearly one third of SIAH1). Such a large N-terminal-truncated mutant SIAH1 might have unknown and unpredictable structural consequences on correct protein folding and thus may yield a possibly incorrect SIAH1 3D structure inappropriate and thus premature for conducting large-scale small molecule inhibitor screens [55-59]. **Thirdly**, several published anti-SIAH inhibition strategies were designed using such truncated SIAH proteins, and several newly identified SIAH small molecule inhibitors were chemically screened and identified by targeting these RING-domain-deleted SIAH mutant proteins [60-64]. For example, an N-terminal truncated SIAH1 (amino acid residues 90–282; i.e., lacking the RING domain) was used as a SIAH inhibitor to reduce tumor growth in melanoma and prostate cancer cells [60]. Another N-terminal truncated SIAH1 (residues 81-282, i.e., missing the RING domain) was used to screen for covalent SIAH inhibitors [61]. A Phyl fragment (1-130 amino acids) was used to inhibit the physical interaction and target degradation of prolyl-hydroxylase (PHD) proteins by SIAH that suppressed mammary tumor growth [64], melanoma metastasis [63], and prostate tumorigenesis and metastasis [62, 65]. Based on our *Drosophila sina* mutagenesis study as presented in here, there is a concern about these SIAH-targeted strategies used in mammalian studies as enlisted above.

Our *Drosophila* mutagenesis screen has revealed three invariable cysteine residues in SINA (C104Y, C73Y, or C76Y substitutions in *sina*^ES10^, *sina*^ES79^, or *sina*^ES473^, respectively). These single point mutations were sufficient to completely suppress *GMR-phyl* rough eye phenotype, block R7 photoreceptor cell fate determination, and impede proper RAS signal transduction in fly eye development. We were the first group to successfully design the point mutations (2 Cys to 2 Ser) in the RING-domain of full-length SIAH1/2 (SIAH1^C41S-C44S^ and SIAH2^C80S-C83S^), named SIAH1^PD^/SIAH2^PD^ [29, 30]. We found that SIAH1^PD^/SIAH2^PD^ functions as potent and long-lasting SIAH inhibitors that could be used to eradicate and abolish oncogenic K-RAS-driven malignant pancreatic and lung cancer in xenograft models [29, 30]. Importantly, our SIAH^PD^ strategy was able to achieve a complete tumor eradication [29, 30]. Following our studies, a similar RING point mutant (H99A/C102A) was proven to be successful to reduce melanoma tumorigenesis and metastasis by both HIF-dependent and -independent mechanisms [63]. In contrast, the SIAH-truncation strategy reduced tumor growth but failed to achieve tumor eradication [62, 64, 65]. Guided by the molecular insights and core principles learned from this unbiased *Drosophila* SINA mutagenesis study, we aim to use the new SINA/SIAH point mutations to re-generate X-ray crystallography as well as design more precise, targeted, and long-lasting SIAH inhibitors to control and eradicate the multidrug-resistant and incurable pancreatic, lung, and triple negative breast cancer in preclinical and co-clinical settings.

Sequencing analysis of the rest of the new *sina* mutant alleles has proven to be technically challenging since a majority of them are embryonic and early larval lethal. By using both the FLP-FRT somatic clonal analysis and FLP-FRT-DFS-OVD germline clonal strategy, we were unable to generate homozygous viable mutant clones, and mutant embryos/eggs after multiple attempts using several “null” *sina* mutant alleles. These results suggested that it is likely that *sina* null alleles are severely growth compromised, embryonic lethal or cell lethal, as compared to its twin clones (marked by bright green cells as labeled by GFP) (**Supplemental Figure S1**). To overcome the cell-lethal and embryonic-lethal phenotypes of *sina* null alleles, we focused on several *sina* mutant alleles that are homozygous viable at 3^rd^ instar larval and early pupal stages (*sina*^Loss-of-function^/*sina*^Loss-of-function^), or heterozygous viable at 3^rd^ instar larval, pupal and adult stages when crossed to the *sina*^3^ mutant allele (*sina*^Loss-of-function^/*sina*^3^). We identified and sequenced several strong *sina*^*EMS*^ mutant alleles whose single point mutation lies in the three invariable cysteine residues of the RING domain that is evolutionarily conserved across all metazoan species, but is absent from the published X-ray crystal structures of the RING-truncated mammalian SIAH1 proteins, suggesting that the mammalian SIAH studies may be aided by our *Drosophila* SINA study (**Figure 1A**). Thus, a large collection of 28 newly identified *sina* mutant alleles offers a unique and valuable opportunity to (1) identify the most critical amino acids and key functional domains that are essential to reveal SIAH biological function downstream of K-RAS signal transduction in *Drosophila* and mice; (2) reveal SIAH as a key tumor vulnerability in oncogenic K-RAS-driven malignant tumor biology; and (3) design a new and better anti-SIAH-based anti-K-RAS/EGFR/HER2 targeted therapy to control and eradicate chemo-resistant, locally advanced and metastatic tumors in the future.

Our sequencing data revealed point mutations resulting in C104Y, C73Y, or C76Y substitutions in *sina*^ES10^, *sina*^ES79^, or *sina*^ES473^, respectively, and a 10 base pair deletion in *sina*^ES26^ (**Figure 1A** and **Table 1**). The three C to Y amino acid substitutions mutated the invariable cysteine (Cys) amino acids that are responsible for zinc coordination in the RING domain of SINA [31, 32] (**Figure 1A**). Without proper SINA function, activation of upstream RAS/RAF/MAPK signals (e.g. constitutively active RAS^V12^) cannot be transmitted in R7 neurons, supporting the notion that SINA functions as a “gatekeeper” critical for transmitting the active RAS signal for neuronal cell fate determination (**Figure 1**). Ectopic expression of *Drosophila* SINA^Wt/PD^ or human SIAH1/2^WT/PD^ using eye-specific drivers results in a rough eye phenotype, disrupting normal photoreceptor cell development and ommatidial assembly (**Figures 2 and 3**). The phenotypes included the missing R7 photoreceptor cells, neuro-degenerative phenotypes in the eye, and defective peripheral nervous system (PNS) development (**Figures 2, 3, and 4**). From our SINA mutagenesis study, the identification of these three invariable Cysteines that control SINA activity, and characterizing *sina* mutant phenotypes in *Drosophila* photoreceptor cell fate determination are crucial to understand SINA/SIAH biology in the context of tumor-driven K-RAS/EGFR/HER2 activation in human cancer. In the future, we will combine the power of a complementary and combinational approach to harness the synergy of *Drosophila* genetics, developmental biology, evolutionary biology, and human cancer biology in better designing the next-generation potent SIAH inhibitors as a new anti-SIAH-based anti-K-RAS/EGFR/HER2 targeted therapy to control and eradicate chemo-resistant, relapsed, locally advanced, and metastatic human cancer. The loss of focal adhesions, cell junctions, and cell morphology changes in response to *sina*^loss of function^ EMS mutations (*sina*^ES10^, *sina*^ES26^, *sina*^ES79^, and *sina*^3^), or SINA^PD^/SIAH^PD^ blockade in the developing larval salivary gland may become pertinent to human cancer biology as we design a synergistic and combinational approach of targeting SIAH and SIAH-interacting proteins in hopes of controlling and eradicating undruggable and incurable oncogenic K-RAS-driven malignant tumors in the future.

*Drosophila* is a powerful, fast, and elegant *in vivo* genetic system to study human SIAH biology. We can use *Drosophila* eye development to rapidly test, validate, and screen for small anti-SIAH peptides that will successfully block RAS-dependent signaling processes in *Drosophila* eye development. This *Drosophila*-centered study highlights that not only are SINA family proteins evolutionarily conserved, but that human SIAH and *Drosophila* SINA are functionally conserved in *Drosophila* neuronal cell development. Due to the extraordinary high degree of evolutionary conservation in this medicinally important family of SINA/SIAH E3 ligases, it is expected that the biological functions of *Drosophila* SINA and human SIAH1/SIAH2 are likely to be functionally conserved or interchangeable (**Figures 2, 3 and 4**). The tools developed in this study, especially the wild type and proteolytic-deficient (PD) human SIAH transgenic flies, will be useful and synergistic in our search for better SIAH inhibitors by conducting rapid SINA/SIAH functional validations in *Drosophila* eye development in support of mammalian SIAH-based human cancer studies in the future.

SIAH is a major tumor vulnerability and the most downstream gatekeeper of the oncogenic RAS/EGFR/HER2 signal transduction cascade [28-32]. Hence, the mechanistic insights of SINA/SIAH function gained from *Drosophila* genetic and mutagenesis studies will be easily translational, clinically relevant, and directly applicable to human cancer biology [66-74]. As we design better SIAH inhibitors to effectively shut down RAS-dependent signaling transmission *in vivo*, we aim to use the *Drosophila* eye, PNS, and salivary gland development as a model system and a rapid drug screening platform to test lead compounds and validate the efficacy of the next-generation anti-SIAH-based anti-K-RAS targeted gene therapy. The complementary approach and synergistic strategy of incorporating this model organism, *Drosophila melanogaster*, into human cancer biology and drug screening pipeline may aid and accelerate our clinical efforts to control and eradicate “undruggable” oncogenic K-RAS/EGFR/HER2-driven malignant cancers in the future.

## Abbreviations

DmSINA^PD^: *Drosophila* SINA^PD^,
DPP: decapentaplegic,
EGFR: epidermal growth factor receptor,
EMS: ethyl methanesulfonate,
ERK: extracellular signal-regulated kinases,
FLP: recombinase that recognizes FRT site,
GAL4: yeast DNA-binding transcriptional activator,
GFP: green fluorescent protein,
GMR: Glass multiple reporter,
hSIAH1^PD^: human SIAH1^PD^,
hSIAH2^PD^: human SIAH2^PD^,
ORF: open reading frame,
PNS: peripheral nervous system,
PD: proteolysis deficient,
RING: really interesting new gene,
SEM: scanning electron microscopy,
SEV: sevenless,
SIAH: human homologs of SINA,
SINA: seven in absentia,
SINA/SIAH inhibitor: SINA^PD^/SIAH^PD^,
UAS: yeast upstream activator sequence,
WT: wild type

## Author contributions

REVS is the 1st author who wrote the 1^st^ draft of this manuscript. AHT is the principal investigator who designed, conducted, and supervised this study. REVS, YC and AHT are involved in the data acquisition, RT-PCR, sequencing analysis, IF staining, confocal imaging capture and graphic design of the figures. REVS generated Figures 1A, 1B, 3, 7, and 8. YC generated Figures 5, 6, 7, and 8, and contributed to Figures 1A and 1B. AHT generated Figures 1A, 1C, 2, and 4. AHT conducted the *GMR-phyl* genetic modifier screen, isolated and identified 28 new *sina* mutant alleles from this modifier screen, established 19 new complementation groups of novel PHYL-interacting proteins, including SINA and EBI, under the advice and guidance of GMR at HHMI and UC Berkeley. AHT designed, cloned, injected, and generated the entire panel of SINA^WT/PD^, SIAH1^WT/PD^, and SIAH2^WT/PD^ transgenes and transgenic flies. AHT conducted the retina section and generated electron microscopy (EM) images of eye and notum phenotypes. AHT designed the figures and wrote the manuscript with the help and input from all team members, especially REVS. AHT is responsible for data management, quality control, statistical analysis, and data interpretation in support of this study.

## Acknowledgements

Correspondence should be addressed to A.H.T. The authors thank Dr. Gerald M. Rubin for his brilliant creativity, high standards, high-quality training, great role modeling, and beautiful scientific guidance in establishing *Drosophila* RAS signaling transduction cascades by conducting elegant genetic screens, and for supporting the *GMR-phyl* F1 modifier screen. The authors thank our colleagues in the Rubin laboratory at UC Berkeley for helpful discussions throughout the course of this work. The authors thank Drs. Jeffery L. Platt, Heidi Nelson, Gloria M. Petersen, Edward B. Leof, the Mayo Clinic Surgery Leadership, and our colleagues at the Xenotransplantation Biology for supporting our *Drosophila* work at the Mayo Clinic. The authors thank Drs. Edward Johnson, John Semmes, Julie Kerry, the EVMS Presidents (Dr. Richard Homan and Mr. Harry Lester), our colleagues at the Department of Microbiology and Molecular Cell Biology and Leroy T. Canoles Jr. Cancer Research Center for supporting our *Drosophila* work at EVMS. The authors thank Dr. Atique Ahmed for his PCR expertise in assisting us with the initial RT-PCR analyses on the heterozygous *sina*^EMS^/*sina*^3^ and *sina*^X-ray^/*sina*^3^ mutant alleles. The authors thank Dr. Minglei Bian for his microscopy expertise in assisting us with confocal imaging capturing and data processing. The authors are grateful to Drs. Rebecca L. Schmidt, Edward B. Leof, and Jeffrey L. Platt for their critical reading of this manuscript and their valuable comments and expert critiques.

## Materials and Methods

### *Drosophila* genetics and mutagenesis

Fly husbandry and genetic mutagenesis were carried out according to standard protocols and procedures. To identify new mutations in the known and novel PHYL-interacting proteins, we conducted an F1 genetic modifier screen in *Drosophila* to isolate dominant enhancers and suppressors of a *GMR-phyl* rough eye phenotype. Isogenic white eye (*w*^-^) male flies were mutagenized either by X-ray irradiation (4000 rad) or ethyl methanesulfonate (EMS) chemical mutagenesis (25 mM). Mutagenized males were then mated *en masse* with virgin females carrying *GMR-phyl* transgenes on the balancer chromosomes, CyO-*GMR-phyl/Adv* or TM3-*GMR-phyl*/*e, ftz, ry*. The F_1_ progeny were scored under a dissection microscope to identify the dominant modifiers of the *GMR-phyl* rough phenotypes. The putative enhancers and suppressors were backcrossed to the CyO-*GMR-phyl or* TM3-*GMR-phyl* screening chromosomes to establish proper chromosomal linkage, balanced them on I, II or III chromosomes, and demonstrate the unambiguous germline transmission of these newly identified PHYL modifiers. Different complementation groups were established by performing genetic complementation tests as described [41, 75-78].

### Transgenic flies carrying *Drosophila* SINA^WT/PD^, human SIAH1^WT/PD^ or SIAH2 ^WT/PD^ genes

Multiple UAS transgenic lines carrying wild type (WT) or proteolytic deficient (PD) variants of full-length *Drosophila* SINA, human SIAH1, or human SIAH2 were established using P-element mediated transformation [79, 80]. To generate SINA/SIAH proteolytic deficient (PD) point mutations, two cysteine amino acids in the zinc-coordinating RING-domain of these E3 ligases were mutagenized as previously described, *Drosophila* SINA (SINA^C73S-C76S^) and human SIAH1/2 protein (SIAH1^C41S-C44S^ and SIAH2^C80S-C83S^) [29, 30].

### RT-PCR and Sequencing from *sina*^*EMS*^ mutant alleles

mRNA was isolated from *sina*^*EMS*^ mutant larvae using RNeasy Mini Kit according to manufacturer’s instructions (Qiagen. Germantown, MD). cDNA synthesis was carried out using AMV First Strand cDNA Synthesis Kit following manufacturer’s instructions (New England BioLabs. Ipswich, MA). PCR amplification was performed using Expand High Fidelity PCR System (Roche. Indianapolis, IN). All primers were purchased from Integrated DNA Technologies (Coralville, IA). The forward and reverse primers for PCR amplification of *sina* coding region, corresponding to the start codon and stop codon, were 5’-ATGTCCAATAAAATCAACCCGAAGCG-3’ and 5’-TTAGACCAGAGATATGGTCACGTTAA-3’, respectively. Ribosomal protein *rp49* mRNA was used as an internal control with forward and reverse primers of 5’-ATACAGGCCCAAGATCGTGA-3’ and 5’-GTGTATTCCGACCACGTTACA-3’, respectively. Sequencing was performed using BigDye terminator kit (v3.1, Life Technologies) and analyzed using an automated sequencer ABI Primer 3130 Genetic Analyzer (Applied Biosystems) according to manufacturer’s instructions.

### Immunofluorescent (IF) staining of imaginal discs and salivary glands in homozygous *sina*^*EMS*^ mutant animals

Immunohistochemical staining of imaginal discs was performed as described [81]. The eye imaginal discs and salivary glands from wandering third instar larvae were dissected and stained as previously described. Briefly, tissues were dissected in 1x phosphate buffered saline (PBS) solution and fixed in 4% paraformaldehyde in PBS for 20 min at room temperature. Eye discs were permeabilized in 0.3% Triton X-100 and 0.3% sodium deoxycholate in 1x PBS. Tissues were blocked in prehybridization buffer (0.3% Triton X-100, 5% Normal Goat Serum, 1% Bovine Serum Albumin in 1x PBS) for 1 h at room temperature. Eye discs were labeled with the following primary antibodies for immunohistochemistry: rat anti-ELAV monoclonal antibody (Developmental Studies Hybridoma Bank, 7E8A10, 1:5 dilution, Iowa City, IA), mouse anti-CUT (Developmental Studies Hybridoma Bank, 2B10, 1:5 dilution, Iowa City, IA). Salivary glands were labeled with the following primary antibodies for immunohistochemistry: rat anti-DE-Cadherin monoclonal antibody (Developmental Studies Hybridoma Bank, DCAD2, 1:20 dilution, Iowa City, IA), mouse anti-Armadillo monoclonal antibody (Developmental Studies Hybridoma Bank, N2 7A1 Armadillo, 1:40 dilution, Iowa City, IA), and mouse anti-α-tubulin (Sigma, T6199, 1:250 dilution). Primary antibody incubations were performed overnight at 4°C. Samples were washed 3 times in 0.1% Triton X-100 in PBS, 10 min each. Tissues were incubated with FITC or Rhodamine Red-X (RRX) conjugated secondary antibodies for 1 hour at room temperature (Jackson ImmunoResearch, used at 1:250 dilution, West Grove, PA). Salivary glands were incubated with Phalloidin–Tetramethylrhodamine B isothiocyanate (Phalloidin-TRITC, Sigma, P1951, 1:300 from 20 µM stock) for 1 hour at room temperature. Samples were washed 3 times in 0.1% Triton X-100 in PBS, 10 min each. Eye discs and salivary glands were incubated in VectaShield mounting medium (Vector Laboratories, Inc. H-1000. Burlingame, CA) for overnight and then mounted on glass slides with coverslips for imaging capturing by fluorescence compound microscopy and/or Zeiss confocal microscopy. IF images were captured with a Leica DMR compound microscope with a 63x (HCX PL APO 63X/1.32 oil PH 3 CS) objective, a Leica DC500 digital camera, and OpenLab software

### *Drosophila* eye sectioning

Fixation, embedding, and sectioning of the adult eye was carried out as previously described [34, 82, 83]. Briefly, heads were dissected away from the adulty fly and one eye cut out to allow for fixatives to penetrate tissue. Heads were fixed in 1% glutaraldehyde and 1 % OsO4 in phosphate buffer for 30 minutes on ice. The tissues were further incubated with fresh OsO4 on ice for 1 hour. Heads were then dehydrated through successive stepwise incubations in 30%, 50%, 70%, 90%, and finally 100% ethanol. A subsequent wash in propylene oxide for 10 minutes at room temperature was repeated twice. Heads were then incubated in an equal volume of propylene oxide and Durcapan Resin overnight at room temperature. Heads were then fixed in pure resin for 4 hours and baked at 70°C overnight. Heads were sectioned in 4 μm thick slices and mounted onto microscope slides and stained with 1% toluidine blue before imaging.

### Electron microscopy of *Drosophila* adult eyes and notum

Young adult flies were prepared for scanning electron microscopy as previously described [34, 84]. Briefly, flies were anesthetized with CO2 and then dehydrated by sequential incubation in ethanol gradient (25%, 50%, 75%, and 100% twice) for 24 hours at each step. Samples were then dehydrated with hexamethyldisilazane and mounted on SEM stubs with TV tube coat and sputter coated with 25-nm thick platinum coat before viewing.

## Supplemental Figures

**Supplemental Figure S1.**
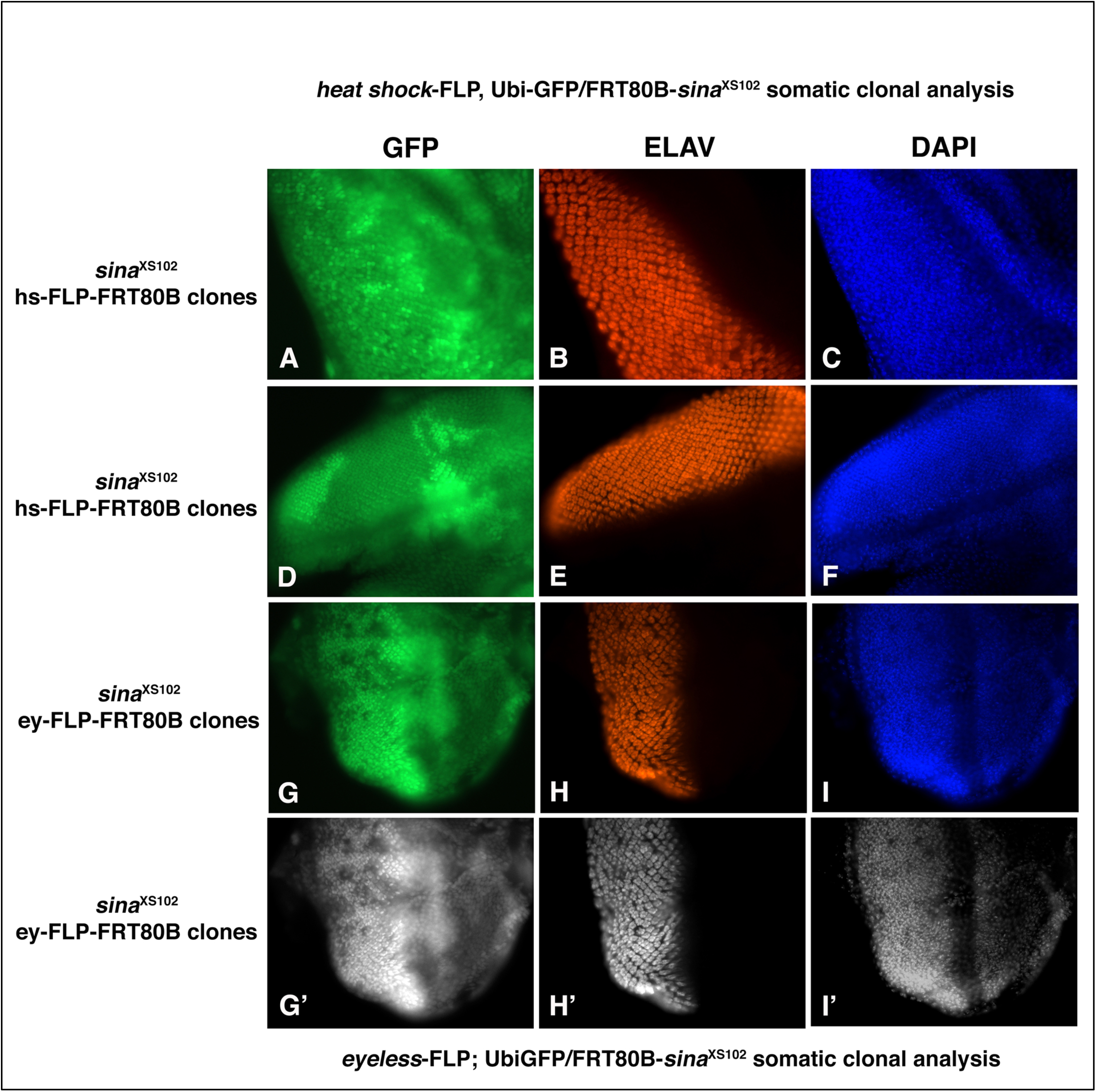
FLP-FRT 80B somatic clonal analysis showed that *sina*^XS102^ *“*null” mutant clones are either severely growth compromised or completely absent as compared to its twin spots (sister clones) in the mosaic analysis. By using FLP-FRT somatic clonal analyses, we generated homozygous *sina* null mutant clones by using both heat shock (hs)-FLP (**A, B, C, D, E** and **F**) and eyeless (ey)-FLP (**G, H, I, G’, H’** and **I’**) to induce somatic clones of *sina* “null” mutant alleles in parallel. Here, *sina*^XS102^ FLP-FRT somatic clones were shown as one example. The FLP-FRT technology was working properly since its twin spots (sister clones), the bright green cells, i.e., double GFP-positive wildtype cells, were abundantly present under the control of either heat shock (hs)-FLP (**A** and **D**) or eyeless (ey)-FLP (**G** and **G’**). In contrast, the *sina* mutant clones, i.e., the GFP-negative cells, were largely missing or severely growth compromised (**B, E, H** and **H**’). These results suggested that *sina* complete loss of function is likely to be cell lethal, as Hoechst staining was also absent in the *sina*^XS102^ mutant clones (GFP-negative cells) (**C, F, I** and **I’**). Two identical sets were shown: **G, H** and **I** are in RGB color, **G’, H’** and **I’** are in black and white color to show the missing cells (black holes) without proper SINA function.

**Supplemental Figure S2.**
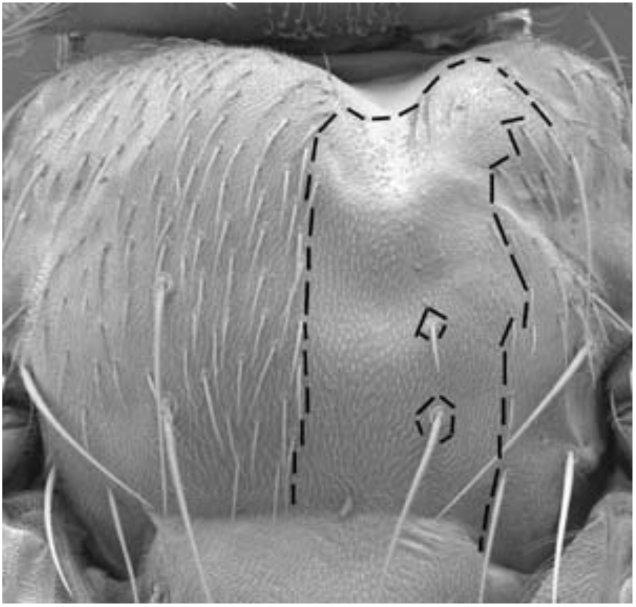
FLP-FRT notum clonal phenotype of a partial loss-of-function alleles of *sina*^*ES26*^. The dashes outline a *sina*^*ES26*^ clone on the notum. This allele is stronger than *sina*^3^.

**Supplemental Figure S3.**
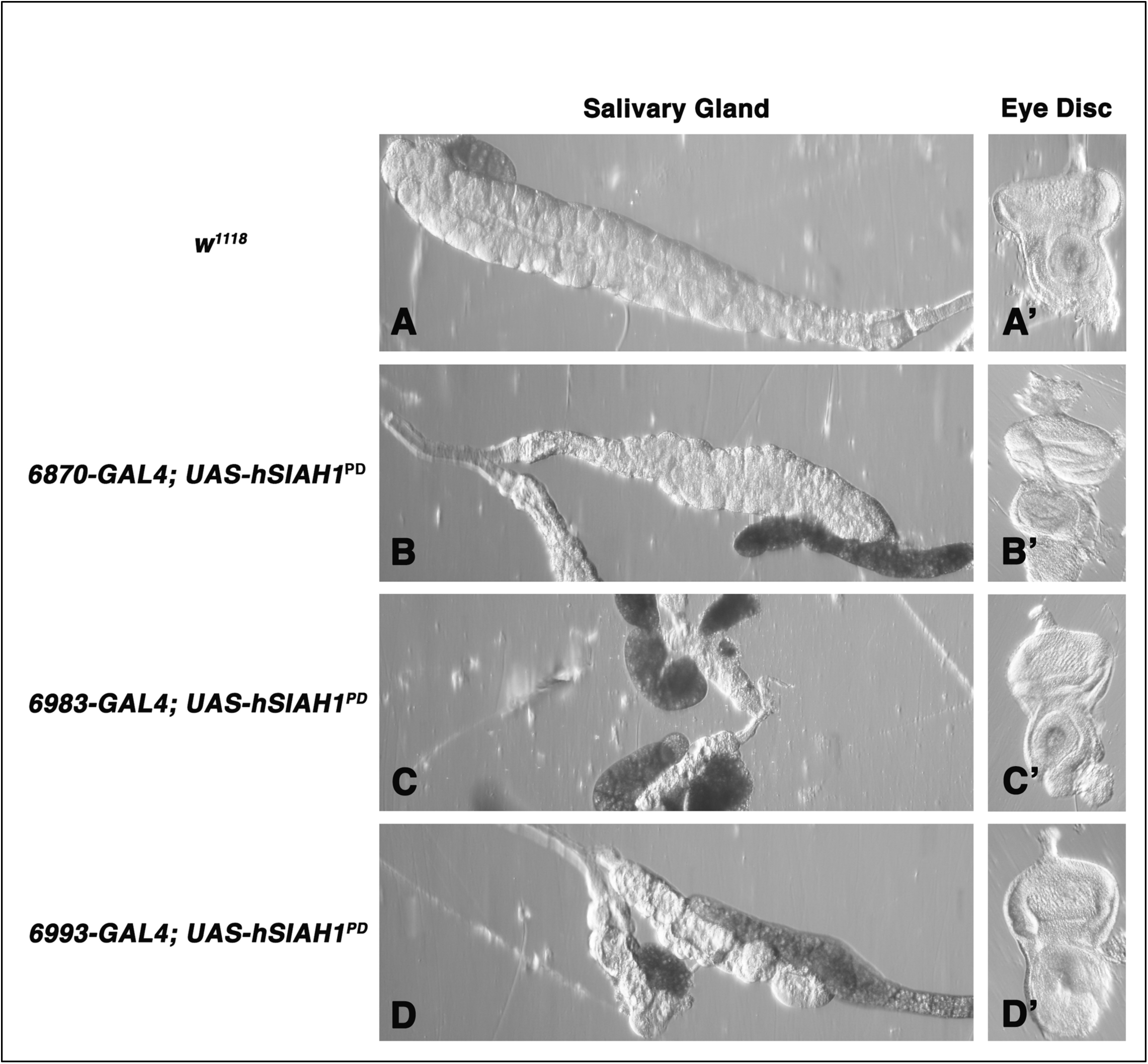
Ectopic expression of human SIAH1 inhibitor, SIAH1^PD^, under the control of three salivary gland-specific GAL4 drivers resulted in cellular defects, junctional disorganization, or tissue atrophy in the larval salivary gland. The normal epithelial cells display highly organized cell junction and hexagon epithelial cell morphology in the wildtype type larval salivary glands under a dissection microscope (**A**). Ectopic expression of SIAH1^PD^ under the control of three salivary gland (SG)-specific GAL4-drivers resulted in cellular defects, junctional disorganization, or tissue atrophy in the larval salivary gland. *6870-GAL4* is a weak SG-driver, *6983-GAL4* is a strong SG-driver, and *6993-GAL4* is an intermediate SG-driver. Ectopic expression of SIAH1^PD^ under the control of the weak *6870-GAL4* resulted in cellular defects, salivary gland disorganization, and mild tissue atrophy (**B**). Ectopic expression of SIAH1^PD^ under the control of the strong *6983-GAL4* resulted in small, disformed, and dying salivary gland with severe tissue atrophy (**C**). Ectopic expression of SIAH1^PD^ under the control of the intermediate *6993-GAL4* resulted in defective, disorganized, and atrophied salivary gland (**D**). Eye discs dissected from the same transgenic animals were used as the internal controls to show the precise timing of the identical developmental stages from which the mutant salivary glands were dissected to examine the tissue morphology and IF staining from the same transgenic animals (**A’, B’, C’** and **D’**).

**Supplemental Figure S4.**
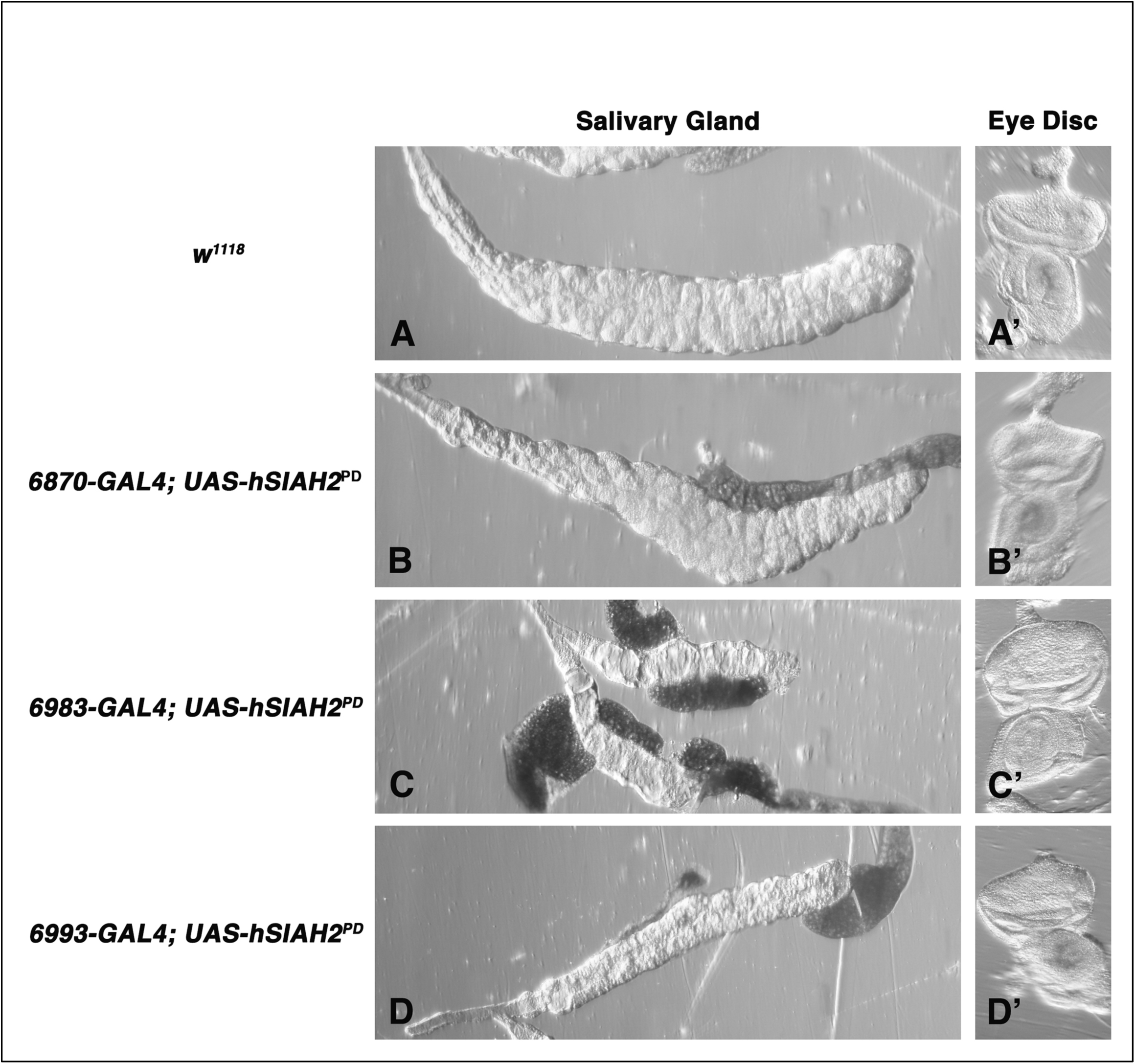
Ectopic expression of human SIAH2 inhibitor, SIAH2^PD^, under the control of three salivary gland-specific GAL4 drivers resulted in cellular defects, junctional disorganization, or tissue atrophy in the larval salivary gland. The normal epithelial cells display highly organized cell junction and hexagon epithelial cell morphology in the wildtype type larval salivary glands under a dissection microscope (**A**). Ectopic expression of SIAH2^PD^ under the control of three salivary gland (SG)-specific GAL4-drivers resulted in cellular defects, junctional disorganization, or tissue atrophy in the larval salivary gland. *6870-GAL4* is a weak SG-driver, *6983-GAL4* is a strong SG-driver, and *6993-GAL4* is an intermediate SG-driver. Ectopic expression of SIAH2^PD^ under the control of the weak *6870-GAL4* resulted in cellular defects, salivary gland disorganization, and mild tissue atrophy (**B**). Ectopic expression of SIAH2^PD^ under the control of the strong *6983-GAL4* resulted in small, disformed, and dying salivary gland with severe tissue atrophy (**C**). Ectopic expression of SIAH2^PD^ under the control of the intermediate *6993-GAL4* resulted in defective, disorganized, and atrophied salivary gland (**D**). Eye discs dissected from the same transgenic animals were used as the internal controls to show the precise timing of the identical developmental stages from which the mutant salivary glands were dissected to examine the tissue morphology and IF staining from the same transgenic animals (**A’, B’, C’** and **D’**).

